# TWIST1 and chromatin regulatory proteins interact to guide neural crest cell differentiation

**DOI:** 10.1101/2020.09.06.285387

**Authors:** Xiaochen Fan, V. Pragathi Masamsetti, Jane Q. J. Sun, Kasper Engholm-Keller, Pierre Osteil, Joshua Studdert, Mark E. Graham, Nicolas Fossat, Patrick P.L. Tam

## Abstract

Protein interaction is critical molecular regulatory activity underlining cellular functions and precise cell fate choices. Using TWIST1 BioID-proximity-labelling and network propagation analyses, we discovered and characterized a TWIST-chromatin regulatory module (TWIST1-CRM) in the neural crest cell (NCC). Combinatorial perturbation of core members of TWIST1-CRM: TWIST1, CHD7, CHD8, and WHSC1 in cell models and mouse embryos revealed that loss of the function of the regulatory module resulted in abnormal specification of NCCs and compromised craniofacial tissue patterning. Our results showed that in the course of cranial neural crest differentiation, phasic activity of TWIST1 and the interacting chromatin regulators promote the choice of NCC fate while suppressing neural stem cell fates, and subsequently enhance ectomesenchyme potential and cell motility. We have revealed the connections between TWIST1 and potential neurocristopathy factors which are functionally interdependent in NCC specification. Moreover, the NCC module participate in the genetic circuit delineating dorsal-ventral patterning of neural progenitors in the neuroepithelium.

## Introduction

The cranial neural crest cell (NCC) lineage originates from the neuroepithelium (Vokes *et al*., 2007; Groves and Labonne, 2014; Mandalos and Remboutsika, 2017) and contributes to the craniofacial tissues in vertebrates (Sauka-Spengler and Bronner-Fraser, 2008) including parts of the craniofacial skeleton, connective tissues, melanocytes, neurons and glia (Kang and Svoboda, 2005; Blentic *et al*., 2008; Ishii *et al*., 2012; Theveneau and Mayor, 2012). The development of these tissues is affected in neurocristopathies, which can be traced to mutations in genetic determinants of NCC specification and differentiation (Etchevers *et al*., 2019). As an example, mutations in transcription factor *TWIST1* in human are associated with craniosynostosis (Ghouzzi *et al*., 2000) and cerebral vasculature defects (Tischfield *et al*., 2017). Phenotypic analyses of *Twist1* conditional knockout mouse revealed that TWIST1 is required in the NCCs for the formation of the facial skeleton, the anterior skull vault, and the patterning of the cranial nerves (Soo *et al*., 2002; Ota *et al*., 2004; Bildsoe *et al*., 2009; Bildsoe *et al*., 2016). To comprehend the mechanistic complexity of NCC development and its implication in a range of diseases, it is essential to collate the compendium of genetic determinants of the NCC lineage and characterize how they act in concert in time and space.

During neuroectoderm development, transcriptional programs are initiated successively in response to morphogen induction to specify neural stem cell (NSC) subdomains along the dorsal-ventral axis in the neuroepithelium (Briscoe *et al*., 2000; Vokes *et al*., 2007; Kutejova *et al*., 2016). NCCs also arise from the neuroepithelium, at the border of the surface ectoderm through pre-epithelial-mesenchymal transition (pre-EMT) which is marked by the activation of *Twist1*, *Tfap2a, Id1, Id2, Zic1, Msx1* and *Msx2* (Baker *et al*., 1997; Mayor *et al*., 1997; Saint-Jeannet *et al*., 1997; Marchant *et al*., 1998; Etchevers *et al*., 2019). In the migratory NCCs, gene activity associated with pre-EMT and NCC specification is replaced by that of EMT and NCC identity (Marchant *et al*., 1998). NCC differentiation progresses in a series of cell fate decisions (Lasrado *et al*., 2017; Soldatov *et al*., 2019). Genetic activities for mutually exclusive cell fates are co-activated in the progenitor population, which is followed by an enhancement of the transcriptional activities that predilect one lineage over the others (Lasrado *et al*., 2017; Soldatov *et al*., 2019). However, more in-depth knowledge of the factors triggering this sequence of events and cell fate bias is presently lacking. Furthermore, it is not clear how NCCs are specified in parallel with other neurogenic cell populations in the neuroepithelium.

*Twist1* expression is initiated during NCC specification and its activity is sustained in migratory NCCs to promote ectomesenchymal fate (Soldatov *et al*., 2019). TWIST1 mediates cell fate choices through functional interactions with other basic-helix-loop-helix (bHLH) factors (Spicer *et al*., 1996; Firulli *et al*., 2005; Connerney *et al*., 2006) in addition to transcription factors SOX9, SOX10 and RUNX2 (Spicer *et al*., 1996; Hamamori *et al*., 1997; Bialek *et al*., 2004; Laursen *et al*., 2007; Gu *et al*., 2012; Vincentz *et al*., 2013). TWIST1 therefore constitutes a unique assembly point to identify the molecular modules necessary for cranial NCC development and determine how they orchestrate the sequence of events in this process.

To decipher the molecular context of TWIST1 activity and identify functional modules, we generated the first TWIST1 protein interactome in NCCs. Leveraging the proximity-dependent biotin identification (BioID) methodology, we captured TWIST1 interactions in the native cellular environment including previously intractable transient low-abundance events which feature interactions between transcription regulators (Roux *et al*., 2012; Kim and Roux, 2016). Integrating prior knowledge of protein associations and applying network propagation analysis (Cowen *et al*., 2017), we uncovered modules of highly connected interactors as potent NCC regulators. Among the top-ranked candidates were histone modifiers and chromatin remodelers that constitute the functional chromatin regulatory module (TWIST1-CRM) in NCC. Genome occupancy, gene expression, and combinatorial perturbation studies of high-ranked members of the TWIST1-CRM during neurogenic differentiation *in vitro* and in embryos revealed their necessity in NCC specification and acquisition of ectomesenchyme potential. This study also identified the concurrent activation and cross-repression of the molecular machinery that governs the choice of cell fate between neural crest and neurogenic cell lineage.

## Results

### Deciphering the TWIST1 protein interactome in cranial NCCs using BioID

The protein interactome of TWIST1 was characterized using the BioID technique which allows for the identification of interactors in their native cellular environment (Figure 1A). We performed the experiment in cranial NCC cell line O9-1 (Ishii *et al*., 2012) transfected with TWIST1-BirA* (TWIST1 fused to the BirA* biotin ligase). In the transfected cells, biotinylated proteins were predominantly localized in the nucleus (Figure S1A, B; (Singh and Gramolini, 2009)). The profile of TWIST1-BirA* biotinylated proteins were different from that of biotinylated proteins captured by GFP-BirA* (Figure S2A). Western blot analysis detected TCF4, a known dimerization partner of TWIST1, among the TWIST1-BirA* biotinylated proteins but not in the control group (Figure S2A). These findings demonstrated the utility and specificity of the BioID technology to identify TWIST1 interacting proteins.

**Figure 1.**
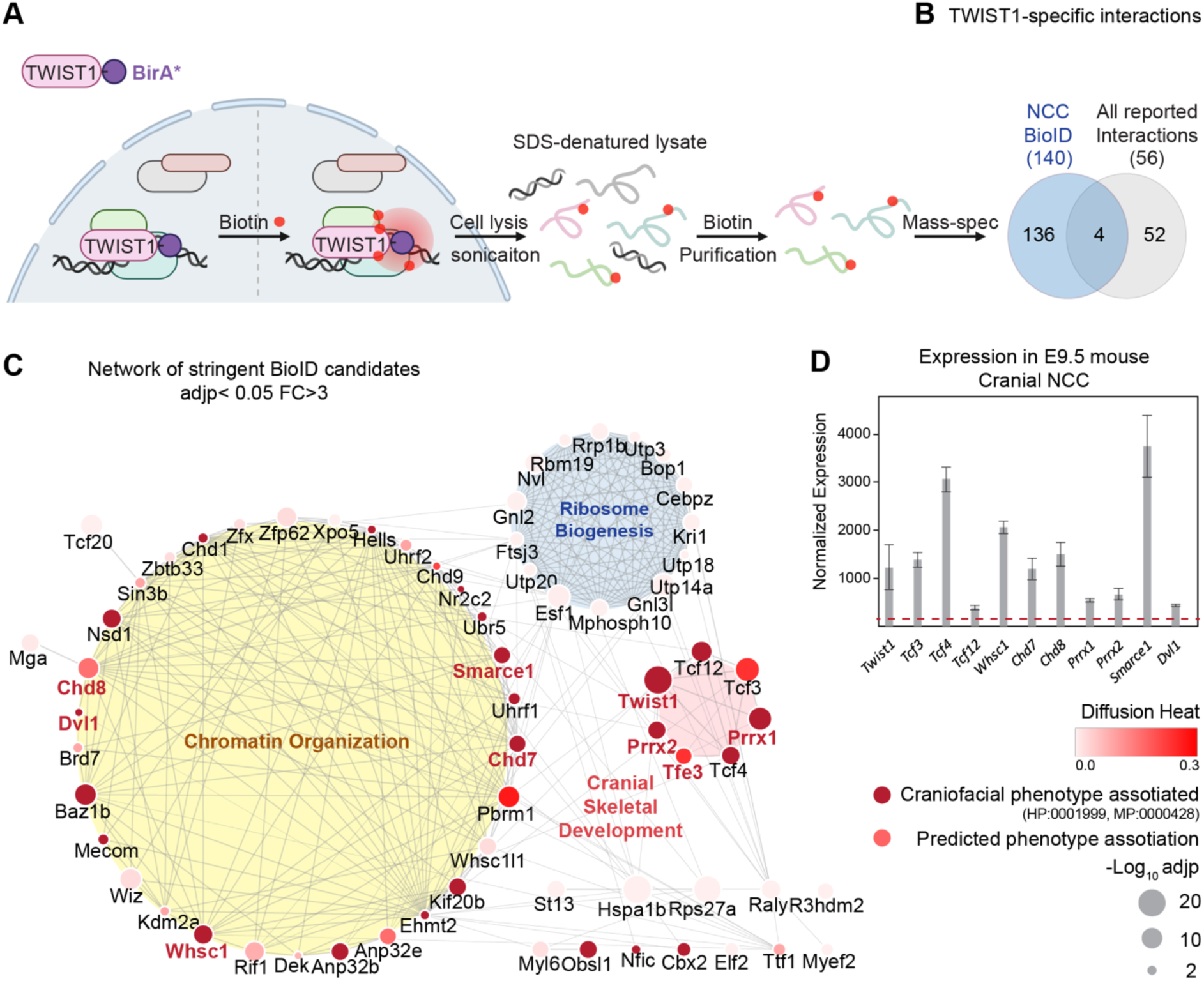
TWIST1 interactome in cranial NCCs revealed using BioID and network propagation. **A.** BioID procedure to identify TWIST1-interacting partners in neural crest stem cells (NCCs). TWIST1-BirA* (TWIST1 fused to the BirA* biotin ligase) labeled the proteins partners within the 10-nm proximity in live cells. Following cell lysis and sonication, streptavidin beads were used to capture denatured biotin-labeled proteins, which were purified and processed for mass spectrometry analysis. **B.** TWIST1-specific interaction candidates identified by BioID mass-spectrometry analysis in NCC cell line (P < 0.05; Fold-change > 3; PSM# > 2) overlap with all reported TWIST1 interactions on the Agile Protein Interactomes DataServer (APID) (Alonso-Lopez *et al*., 2019). **C.** Networks constructed from stringent TWIST1-specific interaction at a significant threshold of adjusted P-value (adjp) < 0.05 and Fold-change > 3. Unconnected nodes were removed. Top GO terms for proteins from three different clusters are shown. Node size = −Log10 (adjp). Genes associated with human and mouse facial malformation (HP:0001999, MP:0000428) were used as seeds (dark red) for heat diffusion through network neighbors. Node color represents the heat diffusion score. **D.** Expression of candidate interactor genes in cranial neural crest from E9.5 mouse embryos; data were derived from published transcriptome dataset (Fan *et al*., 2016). Each bar represents mean expression +/− SE of 3 biological replicates. All genes shown are expressed at level above the microarray detection threshold (red dashed line).

We characterized all the proteins biotinylated by TWIST1-BirA* and GFP-BirA* followed by streptavidin purification using liquid chromatography combined with tandem mass spectrometry (LC-MS/MS) (Table S1). Differential binding analysis of TWIST1 using sum-normalized peptide-spectrum match (PSM) values (Figure S2B, C; see Methods) revealed 140 putative TWIST1 interactors in NCCs (P < 0.05; Fold-change > 3; PSM# > 2; Figure 1B, Table S1). These candidates included 4 of 56 known TWIST1 interactors, including TCF3, TCF4, TCF12 and GLI3 (overlap odds ratio = 18.05, Chi-squared test p-value = 0.0005; Agile Protein Interactomes DataServer [APID]) (Alonso-Lopez *et al*., 2019; Fan *et al*., 2020).

### Network propagation prioritized functional modules and core candidates in TWIST1 interactome

We invoked network propagation analytics to identify functional modules amongst novel TWIST1 BioID-interactors and to prioritize the key NCC regulators (See Methods). Network propagation, which is built on the concept of “guilt-by-association”, is a set of analytics used for gene function prediction and module discovery (Sharan *et al*., 2007; Ideker and Sharan, 2008; Cowen *et al*., 2017). By propagating molecular and phenotypic information through connected neighbors, this approach identified and prioritized relevant functional cluster while eliminating irrelevant ones.

The TWIST1 functional interaction network was constructed by integrating the association probability matrix of the BioID candidates based on co-expression, protein-interaction, and text mining databases from STING (Singh and Gramolini, 2009; Szklarczyk *et al*., 2015). Markov clustering (MCL) was applied to the matrix for the inference of functional clusters (Figure S2D, Table S2). Additionally, data from a survey of the interaction of 56 transcription factors and 70 unrelated control proteins were used to distinguish the most likely specific interactors from the non-specific and the promiscuous TF interactors (Li *et al*., 2015). Specific TF interactors (red) and potential new interactors (blue; Figure S2D-i) clustered separately from the hubs predominated by non-specific interactors (grey; Figure S2D-ii). The stringency of the screen was enhanced by increasing the statistical threshold (adjusted P-value [adjp] < 0.05) and excluding the clusters formed by non-specific interactors such as those containing heat shock proteins and cytoskeleton components. Gene Ontology analysis revealed major biological activities of proteins in the clusters: chromatin organization, cranial skeletal development, and ribosome biogenesis (Figure 1C; Table S2) (Chen *et al*., 2009).

Heat diffusion was applied to prioritize key regulators of NCC development. The stringent TWIST1 interaction network comprises proteins associated with facial malformation phenotypes in human/mouse (HP:0001999, MP:0000428), that points to a likely role in NCC development. These factors were used as seeds for a heat diffusion simulation to find near-neighbors of the phenotype hot-spots (i.e. additional factors that may share the phenotype) and to determine the hierarchical ranking of their importance (Figure 1C, Table S2). Consistent with the expectation that disease causal factors are highly connected and tend to interact with each other (Jonsson and Bates, 2006), a peak of proteins with high degrees of connectivity emerged among the top diffusion ranked causal factors, most of which are from the chromatin organization module (Figure S2F). TWIST1 and these interacting chromatin regulators were referred to hereafter as the TWIST1-chromatin regulatory module (TWIST1-CRM).

Among the top 30 diffusion ranked BioID candidates, we prioritized 9 for further characterization. These included chromatin regulators that interact with TWIST1 exclusively in NCCs versus 3T3 fibroblasts: the chromodomain helicases CHD7, CHD8, the histone methyltransferase WHSC1 and SMARCE1, a member of the SWI/SNF chromatin remodeling complex (Figure 1C, candidates name in red; Figure S2E, F; Table S3). We also covered other types of proteins, including transcription factors PRRX1, PRRX2, TFE3 and the cytoplasmic phosphoprotein DVL1 (Dishevelled 1). The genes encoding these proteins were found to be co-expressed with *Twist1* in the cranial NCCs of in embryonic day (E) 9.5 mouse embryos (Figure 1D) (Bildsoe *et al*., 2016; Fan *et al*., 2016).

### The chromatin regulators interact with the N-terminus domain of TWIST1

Co-immunoprecipitation (co-IP) assays showed that CHD7, CHD8, PRRX1, PRRX2 and DVL1 could interact with TWIST1 like known interactors TCF3 and TCF4, while TFE3 and SMARCE1 did not show any detectable interaction (Figure 2A). Fluorescent immunostaining demonstrated that these proteins co-localized with TWIST1 in the nucleus (Figure S1C). The exceptions were DVL1 and TFE3, which were localized predominantly in the cytoplasm (Figure S1C). Among these candidates, CHD7 and CHD8 are known to engage in direct domain-specific protein-protein interactions (Batsukh *et al*., 2010). Three sub-regions of CHD7 and CHD8 were tested for interaction with TWIST1 (Figure 2B). For both proteins, the p1 region, which encompasses helicases and chromodomains, showed no detectable interaction with partial or full-length TWIST1. In contrast, the p2 and the p3 regions of CHD7 and CHD8 interacted with full-length TWIST1 as well as with its N-terminal region (Figure 2C). Reciprocally, the interaction was tested with different regions of TWIST1 including the bHLH domain, the WR domain, the C-terminal region and the N-terminal region (Figure 2B). CHD7, CHD8 and WHSC1 interacted preferentially with the TWIST1 N-terminus whereas the TCF dimerization partners interacted specifically with the bHLH domain (Figure 2D). Consistent with the co-IP result, SMARCE1 and TFE3 did not interact with TWIST1. Interestingly, the other known factor that binds the TWIST1 N-terminal region is the histone acetyltransferase CBP/P300 which is also involved in chromatin remodeling (Hamamori *et al*., 1999). These findings demonstrated direct interaction of TWIST1 with a range of epigenetic factors and transcriptional regulators and identified the TWIST1 N-terminal region as the domain of contact.

**Figure 2.**
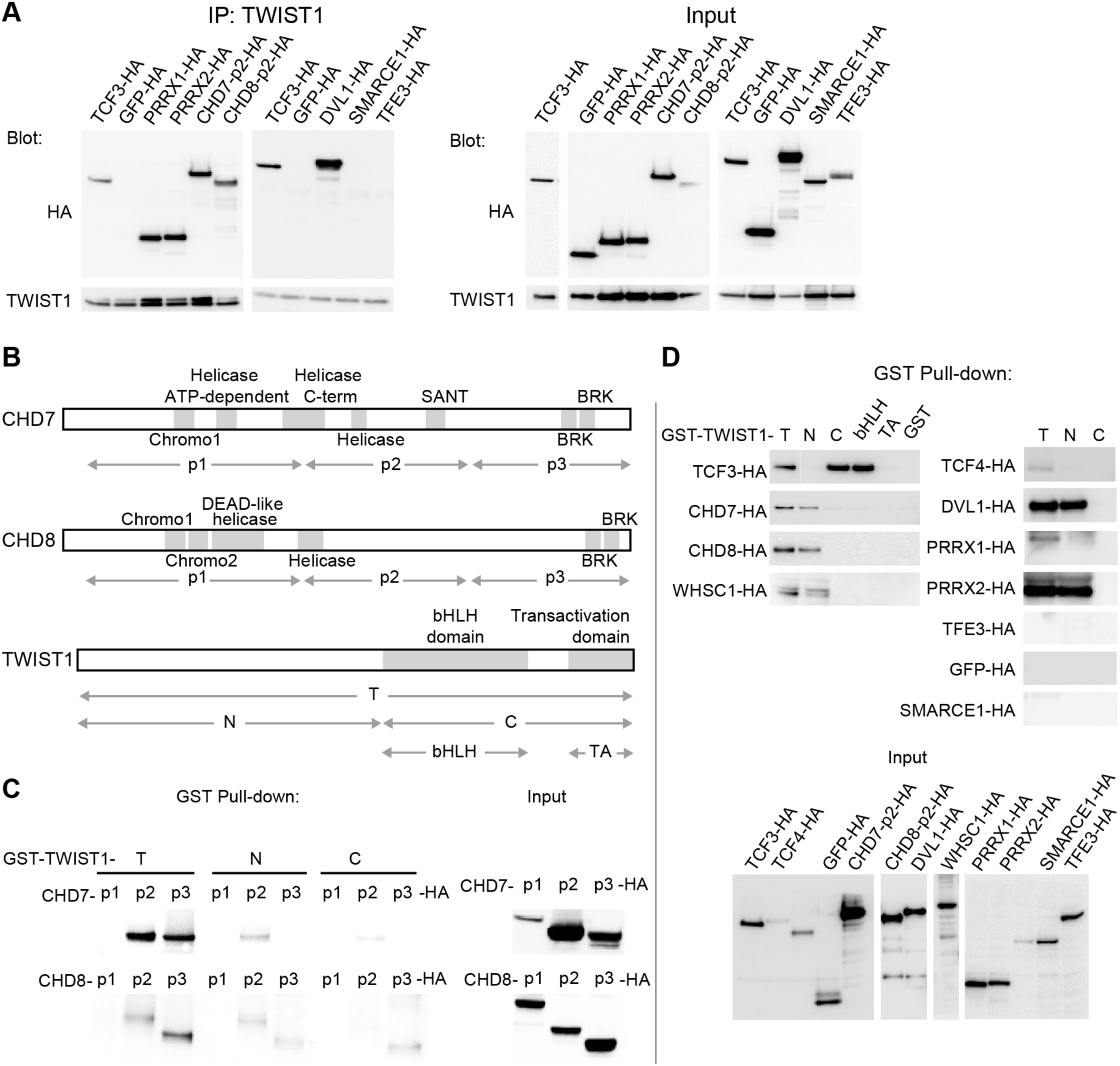
The chromatin regulators interact with the N-terminus domain of TWIST1. **A.** Detection of HA-tagged proteins after immunoprecipitation (IP) of TWIST1 (IP: α-TWIST1) from lysates of cells transfected with constructs expressing TWIST1 (input blot: α-TWIST1) and the HA-tagged proteins partners (input blot: α-HA). **B.** Schematics of CHD7, CHD8 and TWIST1 proteins showing the known domains (grey blocks) and the regions (double arrows) tested in the experiments shown in panels C and D. **C, D.** Western blot analysis of HA-tagged proteins (α-HA antibody) after GST-pulldown with different TWIST1 domains (illustrated in B). Protein expression in the input is displayed separately. T, full-length TWIST1; N, N-terminal region; C, C-terminal region; bHLH, basic helix-loop-helix domain; TA, transactivation domain.

### Genetic interaction of *Twist1* and chromatin regulators in craniofacial morphogenesis

The function of the core components of the TWIST1-CRM was investigated *in vivo* using mouse embryos derived from ESCs that carried single-gene or compound heterozygous mutations of *Twist1* and the chromatin regulators. Mutant ESCs for *Twist1* and the three validated NCC-exclusive chromatin regulatory partners *Chd7, Chd8 and Whsc1* were generated by CRISPR-Cas9 editing (Figure S3A, B) (Ran *et al*., 2013). ESCs of specific genotype (non-fluorescent) were injected into 8-cell host wildtype embryos (expressing fluorescent DsRed.t3) and chimeras were collected at E9.5 or E11.5 (Figure 3A) (Sibbritt *et al*., 2019). Only embryos with predominant contribution of mutant ESCs, indicated by absence or low level of DsRed.t3 fluorescence were analyzed. The majority of embryos derived from single-gene heterozygous ESCs (*Twist1^+/−^, Chd7^+/−^, Chd8^+/−^* and *Whsc1^+/−^*) displayed mild deficiency in the cranial neuroepithelium and focal vascular hemorrhage (Figure 3B). Compound heterozygous embryos (*Twist1^+/−^;Chd7^+/−^, Twist1^+/−^;Chd8^+/−^* and *Twist1^+/−^;Whsc1^+/−^*) displayed more severe craniofacial abnormalities and exencephaly (Figure 3B, C).

**Figure 3.**
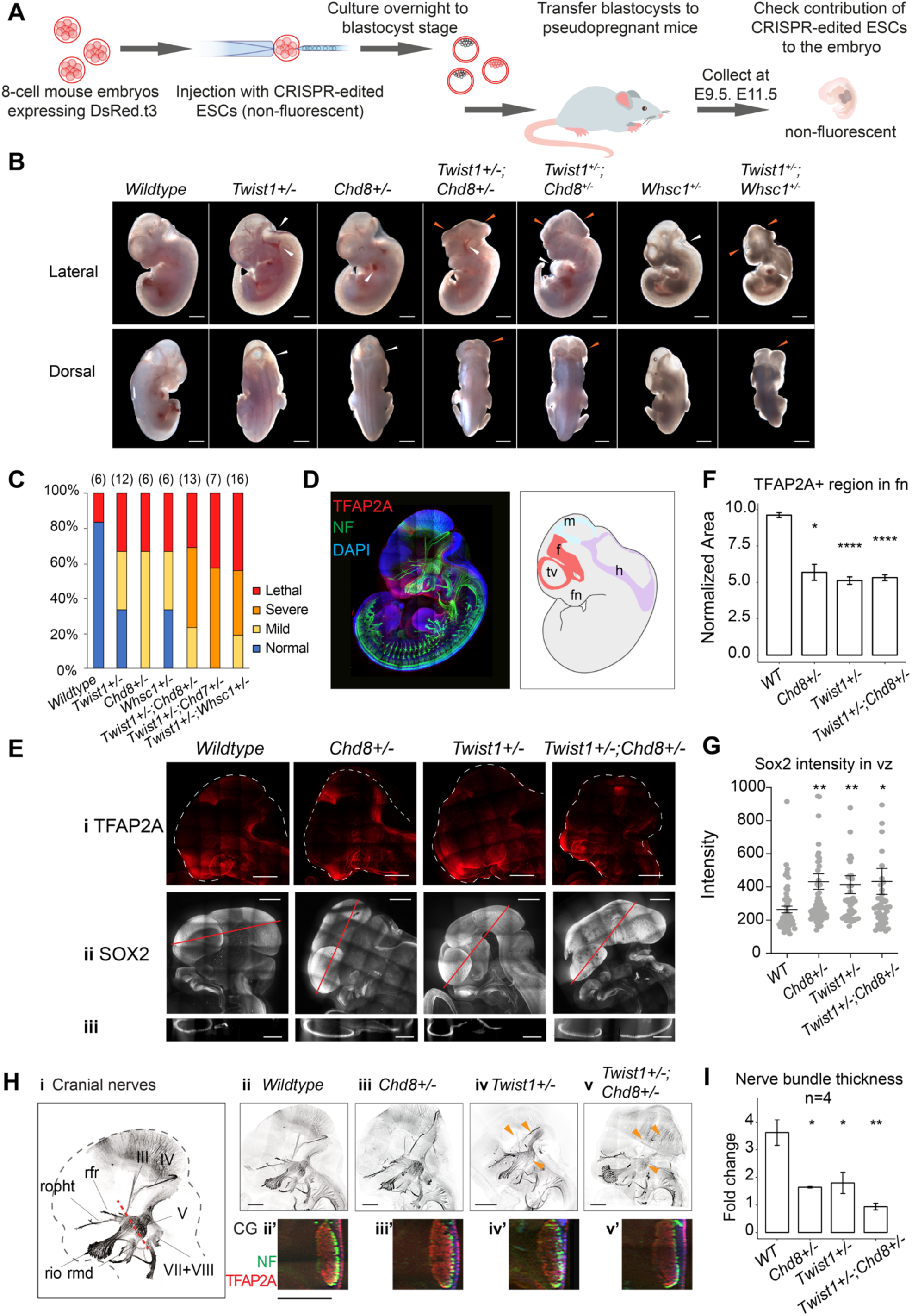
Genetic interaction of Twist1 and chromatin regulators in craniofacial morphogenesis. **A.** Experimental strategy for generating chimeric mouse from WT and mutant ESCs (see Methods). **B.** Lateral and dorsal view of mid-gestation chimeric embryos with predominant ESC contribution (embryo showing low red fluorescence). Genotype of ESC used for injection is indicated. Scale bar: 1mm. Heterozygous embryos of single genes (*Twist1^+/−^, Chd8^+/−^, Whsc1^+/−^*) showed mild defects including hemorrhages and mild neural tube defect (white arrows). Compound heterozygous embryos displayed open neural tube and head malformation (orange arrows, n ≥ 6 for each genotype), in addition to heart defects. **C**. Proportions of normal and malformed embryos (Y-axis) for each genotype (X-axis). Severity of mutant phenotypes was determined based on the incidence of developmental defects in neuroepithelium, midline tissues, heart and vasculature: Normal (no defect); Mild (1-2 defects); Severe (3-4 defects) and early lethality. The number of embryos scored for each genotype is in parentheses. **D.** Whole-mount immunofluorescence of E11.5 chimeras derived from wildtype ESCs, shows the expression of TFAP2A (red) and neurofilament (NF, green) and cell nuclei by DAPI (blue). Schematic on the right shows the neuroepithelium structures: f, forebrain; m, midbrain; h, hindbrain; tv, telencephalic vesicle; fn, frontonasal region. **E. i.** NCC cells, marked by TFAP2A, and neuroepithelial cells, marked by SOX2, are shown in **ii.** sagittal and **iii.** transverse view of the craniofacial region (red dashed line in ii: plane of section). **F.** Quantification of frontal nasal TFAP2A+ tissues (mean normalized area +/− SE) of three different sections for each genotype. **G.** SOX2 intensity (mean +/− SE) in the ventricular zone of three sections for each genotype were quantified using IMARIS. **H. i.** Cranial nerve structures were visualized by immuno-staining of neurofilament (NF). **ii-v** maximum projection of cranial nerves in embryos. Missing or hypoplastic neurites are indicated by arrowheads. **ii’-v’** cross-section of neurofilament bundles at trigeminal ganglion (V). Red dashed line in i: plane of section. Bar: 500 μm; V, trigeminal ganglion; III, IV, VII, VIII; rio, infraorbital nerve of V2; rmd, mandibular nerve; ropht, ophthalmic profundal nerve of V1; rfr, frontal nerve. **I.** Thickness of neural bundle in the trigeminal ganglion was measured by the GFP positive area, normalized against area of the trigeminal ganglion (TFAP2A+). Values plotted represent mean fold change +/− SE. Each condition was compared to *WT*. P-values were computed by one-way ANOVA. *P < 0.05, **P < 0.01, ***P < 0.001, **** P < 0.0001. ns, not significant.

In view of that CHD8 was not previously known to involve in craniofacial development of the mouse embryo, we focused on elucidating the impact of genetic interaction of *Chd8* and *Twist1* on NCC development *in vivo.* While *Chd8*^+/−^ embryos showed incomplete neural tube closure, compound *Twist1*^+/−^*;Chd8*^+/−^ embryos displayed expanded neuroepithelium, a phenotype not observed in the single-gene mutants (Figure 3B, E). The population of NCCs expressing TFAP2α, a TWIST1-independent NCC marker (Brewer *et al*., 2004) was reduced in the frontonasal tissue and the trigeminal ganglion (Figure 3E-i, F). In contrast, SOX2 expression was upregulated in the ventricular zone of the neuroepithelium of mutant chimeras (Figure 3E-ii, iii, G). Furthermore, *Twist1*^+/−^, *Chd8*^+/−^ and *Twist1*^+/−^;*Chd8*^+/−^ embryos displayed different degrees of hypoplasia of the NCC-derived cranial nerves (Figure 3H). Cranial nerves III and IV were absent, and nerve bundle in the trigeminal ganglia showed reduced thickness (Figure 3H, I) most evidently in the *Twist1^+/−^;Chd8^+/−^* compound mutant embryos (Figure 3F-v,v’, I). Altogether, these results suggested that TWIST1 genetically interaction with epigenetic regulators CHD7, CHD8 and WHSC1 to guide the formation of the cranial NCC and downstream tissue genesis *in vivo*.

### Genomic regions co-bound by TWIST1 and chromatin regulators are enriched for active regulatory signatures and neural tube patterning genes

The phenotypic data so far indicate that the combined activity of TWIST1-chromatin regulators might be required for the establishment of NCC identity. To examine whether TWIST1-chromatin regulators are required for NCC specification from the neuroepithelium and to pinpoint its primary molecular function in early neural differentiation, we performed an integrative analysis of ChIP-seq datasets of the candidates. The ChIP-seq dataset for TWIST1 was generated from the ESC-derived neuroepithelial cells (NECs) which are progenitors of NCCs (Figure S4 and Methods). We retrieved published NEC ChIP-seq datasets for CHD7 and CHD8 and the histone modifications and reanalysed the data following the ENCODE pipeline (Consortium, 2012; Sugathan et al., 2014; Ziller et al., 2015) (Figure S4A). Two H3K36me3 ChIP-seq datasets for NECs were included in the analysis on the basis that WHSC1 trimethyl transferase targets several H3 lysine (Morishita *et al*., 2014) and catalyzes H3K36me3 modification *in vivo* (Nimura *et al*., 2009).

Genome-wide co-occupancies of TWIST1, CHD7 and CHD8 showed significant overlap (Fisher’s exact test) and clustered by Jaccard Similarity matrix (Figure 4A). ChIP-seq peaks were correlated with active histone modifications H3K27ac and H3K4me3 but not the inactive H3K27me3, or the WHSC1-associated H3K36me3 modifications (Figure 4A). TWIST1, CHD7 and CHD8 shared a significant number of putative target genes (Figure 4B). TWIST1 shared 63% of target genes with CHD8 (odds ratio = 16.93, Chi-squared test p-value < 2.2e-16) and 18% with CHD7 (odds ratio = 8.26, p-value < 2.2e-16; Figure 4B; Table S4). Compared with genomic regions occupied by no or only one factor, greater percentage of regions with peaks for two or all three factors (TWIST1, CHD7 and CHD8) showed H3K27ac and H3K4me3 signal (Figure 4C). This trend was not observed for the H3K27me3 modification. Similarly, the co-occupied transcription start sites (TSS) showed active chromatin signatures with enrichment of H3K4me3 and H3K27Ac and depletion of H3K27me3 (Ernst *et al*., 2011; Rada-Iglesias *et al*., 2011)(Figure 4D, E). We also did not observe H3K36me3 modifications near the overlapping TSSs, suggesting that WHSC1 may have alternative histone lysine specificity in the NECs.

**Figure 4.**
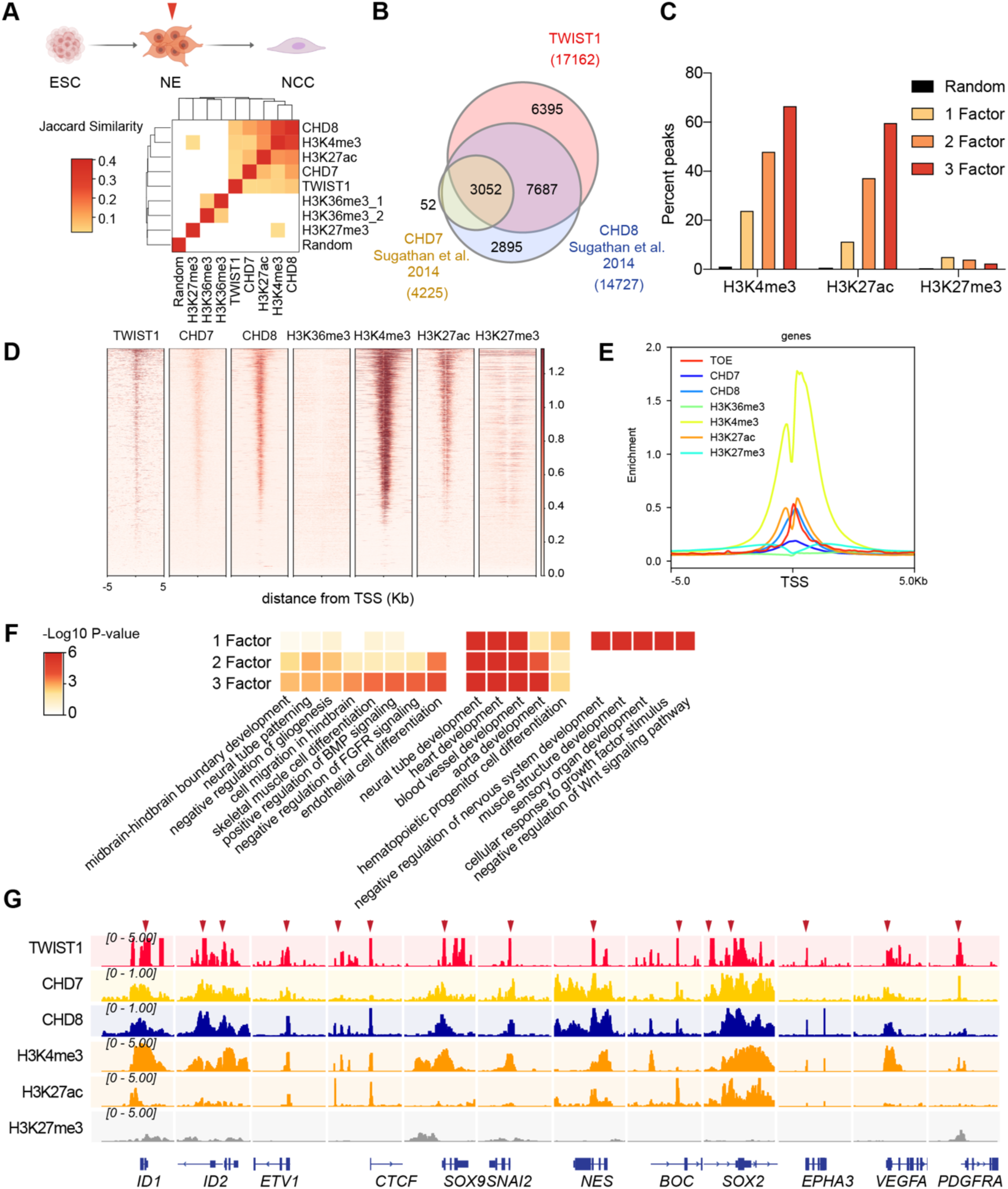
Genomic region showing overlapping binding of TWIST1 and partners are enriched for active regulatory signatures and neural tube patterning genes. **A.** Top panel: Trajectory of ESC differentiation to neuroepithelial cells (NECs) and NCCs. Bottom panel: Jaccard Similarity matrix generated of ChIP-seq data of TWIST1, CHD7, CHD8 and histone modifications from NE cells. The Jaccard correlation is represented by a color scale. White squares indicate no significant correlation (P<0.05, fisher’s exact test) or odds ratio < 10 between the two datasets. **B.** Venn diagram showing overlaps of putative direct target of TWIST1, CHD7, and CHD8, based on ChIP-seq datasets for NE cells (Sugathan *et al*., 2014). **C.** Percent genomic region that is marked by H3K4me3, H3K27ac and H3K27me3 among regions bound by one, two or all three factors among TWIST1, CHD7 and CHD8. Randomized peak regions of similar length (1 kb) were generated for hg38 as a control. **D.** Heatmaps of genomic footprint of protein partners at +/− 5kb from the TSS, based on the ChIP-seq datasets (as in A) and compared with histone marks H3K4me3, H3K27ac and H3K27me3 in human neural progenitor cells (Ziller *et al*., 2015). TSS lanes with no overlapping signals were omitted. **E.** ChIP-seq density profile (rpkm normalized) for all TSS flanking regions shown in D. **F.** Gene Ontology analysis of genomic regulatory regions by annotations of the nearby genes. Regions were grouped by presence of binding site of individual factor (TWIST1, CHD7 and CHD8), or combinatorial binding of 2 or 3 factors. The top non-redundant developmental processes or pathways for combinatorial binding peaks or individual factor binding peaks are shown. P-value cut-off: 0.05. **G.** IGV track (Robinson *et al*., 2011) showing ChIP-peak overlap (red arrows) at common transcriptional target genes in neurogenesis and cell mobility in NCC development. Gene diagrams are indicated (bottom row).

The top Gene Ontology enriched for the co-occupied regulatory regions of 2 or 3 core components included neural tube patterning, cell migration and BMP signaling pathway (Figure 4F). Regions with single factor binding sites were specifically enriched for different sets of ontology such as negative regulation of nervous system and muscle development. Overlapping peaks of the partners were localized within +/− 1 kb of the TSS of common target genes (Figure 4G; Table S4). This integrative analysis revealed that the TWIST1-chromatin regulators shared genomic targets that are harbored in open chromatin regions in the NECs. Therefore, combinatorial binding sites for TWIST1, CHD7 and CHD8 may confer specificity for regulation of patterning genes in the NECs.

### TWIST1 is required for the recruitment of CHD8 to the regulatory region of target genes

To examine whether TWIST1 is necessary to recruit partner proteins to specific regions of co-regulated genes or vice versa, we examined chromatin binding of the endogenous proteins in NECs by ChIP-qPCR analysis (Figure 5A). As CHD8 correlate best with TWIST1 in their ChIP-seq profile surrounding TSS, we analyzed the pattern of recruitment of TWIST1 and CHD8 at the shared peaks near *Sox2, Epha3, Pdgfrα* and *Vegfα* (Figure 4G). One of the peaks near the *Sox2* TSS demonstrated binding by both TWIST1 and CHD8 (Figure 5A, B). In *Twist1^+/−^* or *Chd8^+/−^* NECs, the binding of TWIST1 or CHD8 at the peak were reduced. Interestingly, *Twist1^+/−^* mutation also diminished the binding of CHD8 yet *Chd8^+/−^* mutation did not affect TWIST1 binding (Figure 5A). For *Epha3, Vegfα* and *Pdgfrα*, peaks identified by ChIP-seq with H3K4me3 or H3K27ac modifications were tested (Figure 4G). Partial loss of *Twist1* significantly affected the recruitment of both TWIST1 and CHD8 but again, the loss of CHD8 only affected its own binding (Figure 5C-E). These findings support that TWIST1 binding is a prerequisite for the recruitment of CHD8.

**Figure 5.**
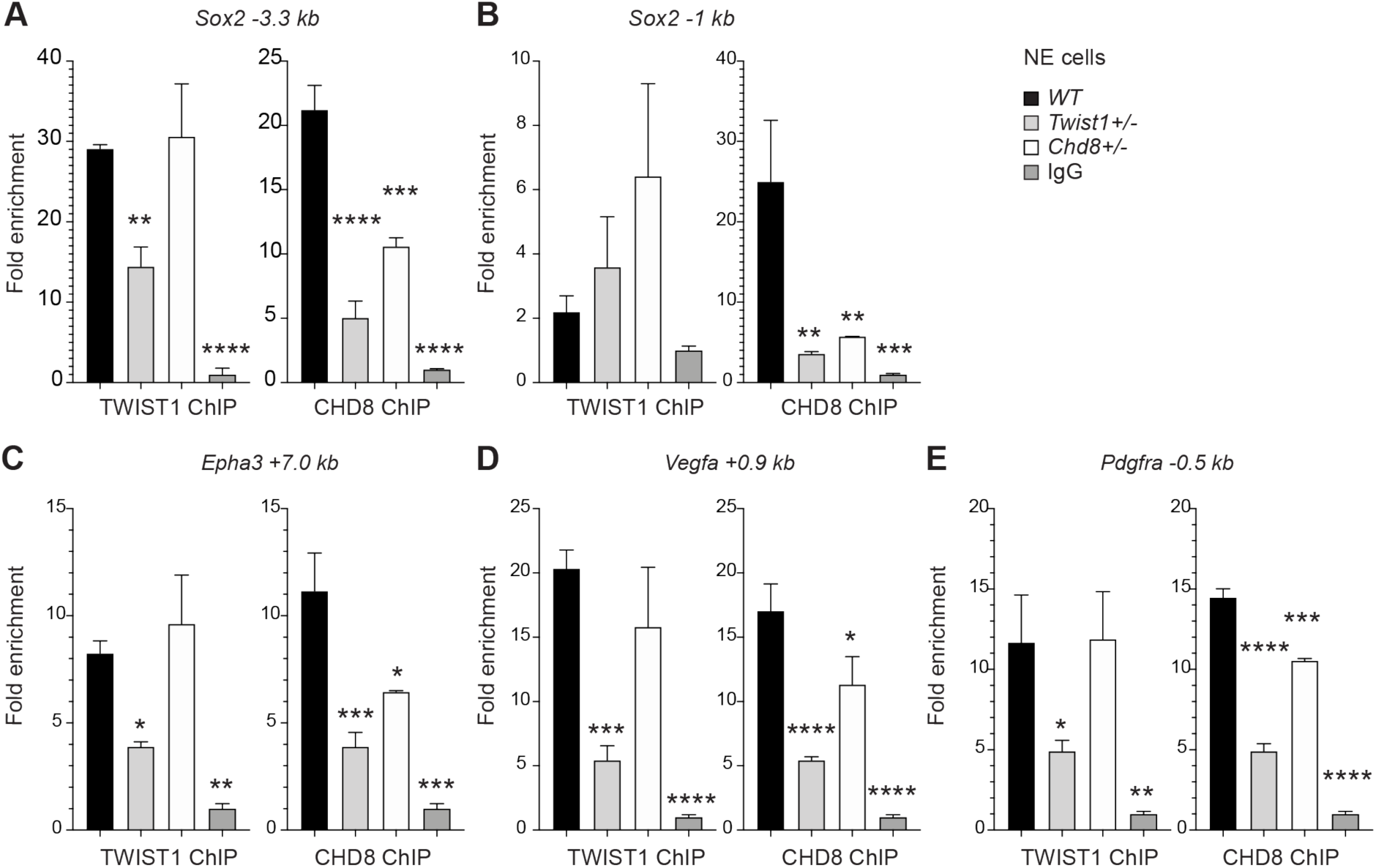
TWIST1 is required for the recruitment of CHD8 to the regulatory region of target genes. Binding of endogenous TWIST1 and CHD8 to overlapping genomic peak regions called by MACS2 (q < 0.05) were assessed by ChIP-qPCR. **A-E.** qPCR quantification of genomic DNA from ChIP of endogenous TWIST1 or CHD8 proteins are shown as mean fold enrichment +/− SE. ChIP experiments using anti-TWIST1 or anti-CHD8 antibodies against endogenous proteins were performed on *wildtype* (*WT)*, *Twist1*^+/−^ and *Chd8*^+/−^ NECs derived from ESC (n = 3, day 3). qPCR results were normalized against signal from non-binding negative control region and displayed as fold change against IgG control. Each condition was compared against *WT* and P-values were generated using one-way ANOVA. *P < 0.05, **P < 0.01, ***P < 0.001, **** P < 0.0001. ns, not significant.

### The TWIST1-chromatin regulators are necessary for cell migration and NCC ectomesenchyme potential

As the TWIST1 and partners were found to regulate cell migration and BMP signaling pathways through target gene binding, we again took a loss-of-function approach and examined the synergic function of TWIST1-chromatin regulatory factors on cell motility in both NECs and NCCs. The migration of NECs out of their colonies was captured by time-lapse imaging and were quantified (see Methods). While *Chd7*^+/−^, *Chd8*^+/−^ and *Whsc1*^+/−^ mutant cells displayed marginally reduced motility, the motility of the *Twist1*^+/−^ cells was compromised and further reduced in *Twist1*^+/−^; *Chd7*^+/−^*, Twist1*^+/−^; *Chd8*^+/−^, and *Twist1*^+/−^; *Whsc1*^+/−^ compound mutant cells (Figure 6A, B).

**Figure 6.**
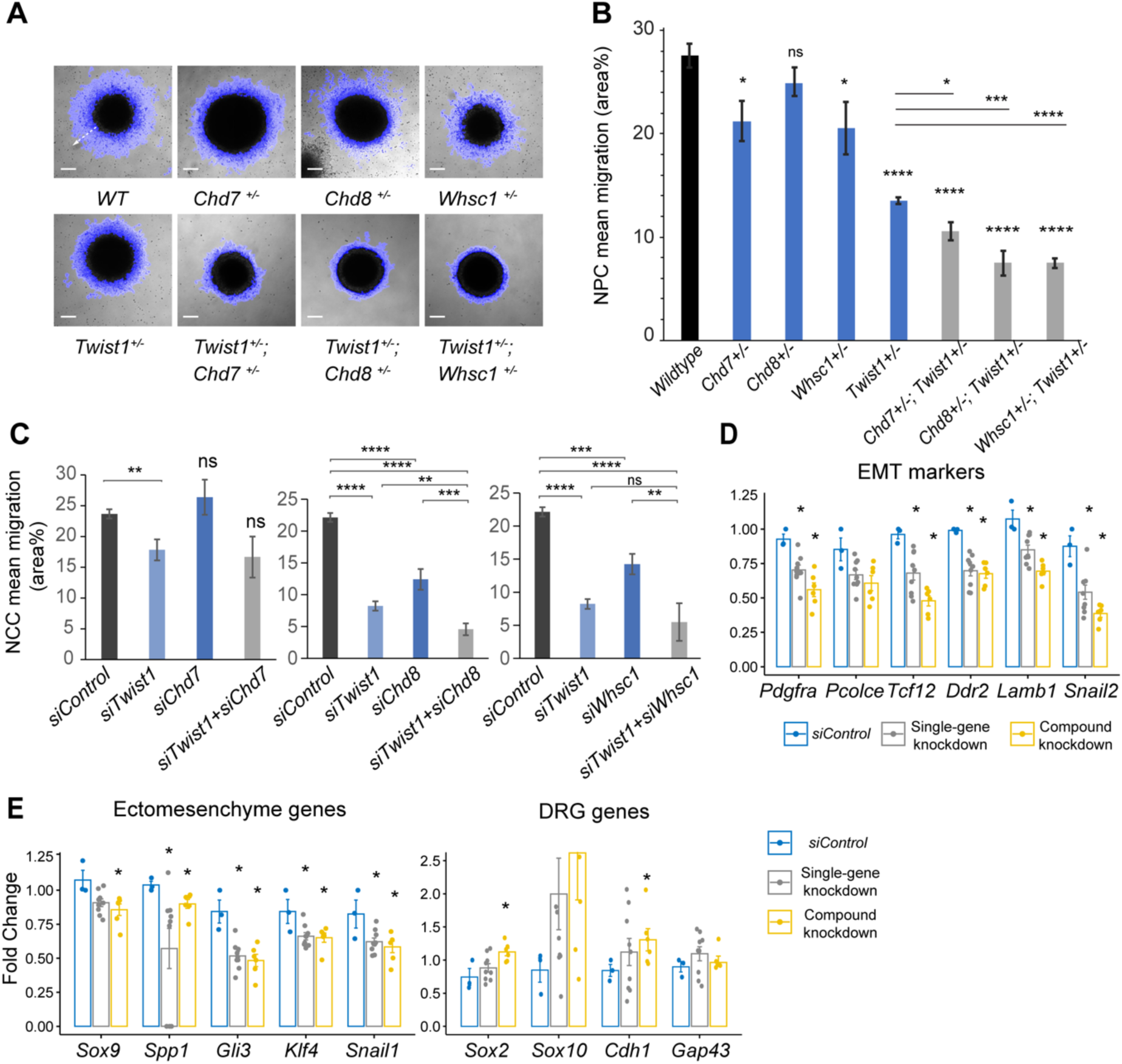
The TWIST1-chromatin regulators are necessary for cell migration and the NCC ectomesenchyme potential. **A.** Dispersion of cells from the colony over 10 hours period *in vitro* (blue halo area). White arrow (shown in wildtype, *WT*) indicates the centrifugal cell movement. Bright-field time-lapse images were captured at set tile regions. Bar = 0.2 mm **B.** Cell migration over 10 hours was quantified from time-lapse imaging data and plotted as mean area % +/− SE for each cell type. n = 5 for each genotype. P-values computed by one-way ANOVA with Holm-sidak post-test**. C.** Results of the scratch assay of O9-1 cells with siRNA knockdowns of *Twist1*, *Chd7, Chd8, Whsc1* and control siRNA. Bright-field images were captured at set tile regions every 15 mins over a 10-hour period. Cell migration was measured as mean area % traversed +/− SE, in triplicate experiments for each genotype. Each condition was compared to *WT*. P-values computed by one-way ANOVA. *P < 0.05, **P < 0.01, ***P < 0.001, **** P < 0.0001. ns, not significant. **D, E.** RT-qPCR quantification of expression of genes associated with EMT, ectomesenchyme, and genes upregulated in the dorsal root ganglia (DRG) vs ectomesenchyme in NCCs. Gene expression is represented as fold change against control +/− SE. Bar diagram shows the expression fold changes in cells treated with *siRNA* individually for *Twist/ Chd8* or *Whsc1* (grey bar) and *siRNA* for *Twist1* in combination with *Chd8* or *Whsc1* (yellow bar). Expression were normalized with the average expression of 3 housekeeping genes (*Gapdh, Tbp, Actb*). Each group was compared to control knockdown treatment. P-values computed by one-way ANOVA. *P < 0.05.

The migratory capacity was also evaluated in O9-1 cells, which are developmentally equivalent to ectomesenchymal NCCs (Ishii *et al*., 2012). NCCs were treated with siRNA to knockdown *Chd7, Chd8* and *Whsc1* activity individually (single-gene knockdown) and in combination with *Twist1-*siRNA (compound knockdown; Figure S3C, see Methods). NCCs treated with *Chd8*-siRNA or *Whsc1-*siRNA but not *Chd7*-siRNA showed impaired motility (relative to control-siRNA treated cells), which was exacerbated by the additional knockdown of *Twist1* (Figure 6C). Impaired motility in *Twist1, Chd8* and *Whsc1* knockdowns was accompanied by reduced expression of EMT genes (*Pdgfrα, Pcolce, Tcf12, Ddr2, Lamb1 and Snai2*) (Figure 6D, S3D) and ectomesenchyme markers (*Sox9, Spp1, Gli3, Klf4, Snai1*), while genes that are enriched in the sensory neurons located in the dorsal root ganglia (Ishii *et al*., 2012) were upregulated (*Sox2, Sox10, Cdh1, Gap43*; Figure 6E). Combined knockdowns had a stronger impact on the expression of the target genes than individual knockdowns for *Twist1, Chd8* and *Whsc1* (Figures 6D, E; S3D). These findings suggest that the acquisition of ectomesenchyme propensity (cell mobility, EMT and mesenchyme differentiation) requires the activity of TWIST1-CHD8/WHSC1.

### TWIST1 and chromatin regulators for cell fate choice in neuroepithelial cells and lineage trajectory of neural crest cells

The genomic and embryo phenotypic data collectively suggest a requirement of TWIST1-chromatin regulators in the establishment of NCC identity in heterogeneous neuroepithelial populations. To understand how TWIST1-chromatin regulators coordinates NCC and other identities during neural differentiation, we studied the module factors during *in vitro* neural differentiation of ESCs. We assessed the lineage propensity of neuroepithelial cells derived from single-gene heterozygous ESCs (*Twist1^+/−^, Chd7^+/−^, Chd8^+/−^* and *Whsc1^+/−^*) and compound heterozygous ESCs (*Twist1^+/−^;Chd7^+/−^, Twist1^+/−^;Chd8^+/−^* and *Twist1^+/−^;Whsc1^+/−^*). ESCs were cultured in neurogenic differentiation media, followed by selection and expansion of NECs (Figure 7A) (Bajpai *et al*., 2010; Varshney *et al*., 2017). Samples were collected at day 0 (ESCs), day 3 and day 12 of differentiation and assessed for the expression of cell markers and ChIP-seq target genes (Table S5). All cell lines progressed in the same developmental trajectory (Figure 7B-i) and generated Nestin-positive rosettes typical of NECs (Figure 7D). Genes were clustered into three groups by patterns of expression: activation, transient activation, and repression (Figure 7B ii, black, red, grey clusters). Notably, *Chd7, Chd8* and *Whsc1* clustered with NCC specifiers that were activated transiently during differentiation (Figure 7B ii, red).

**Figure 7.**
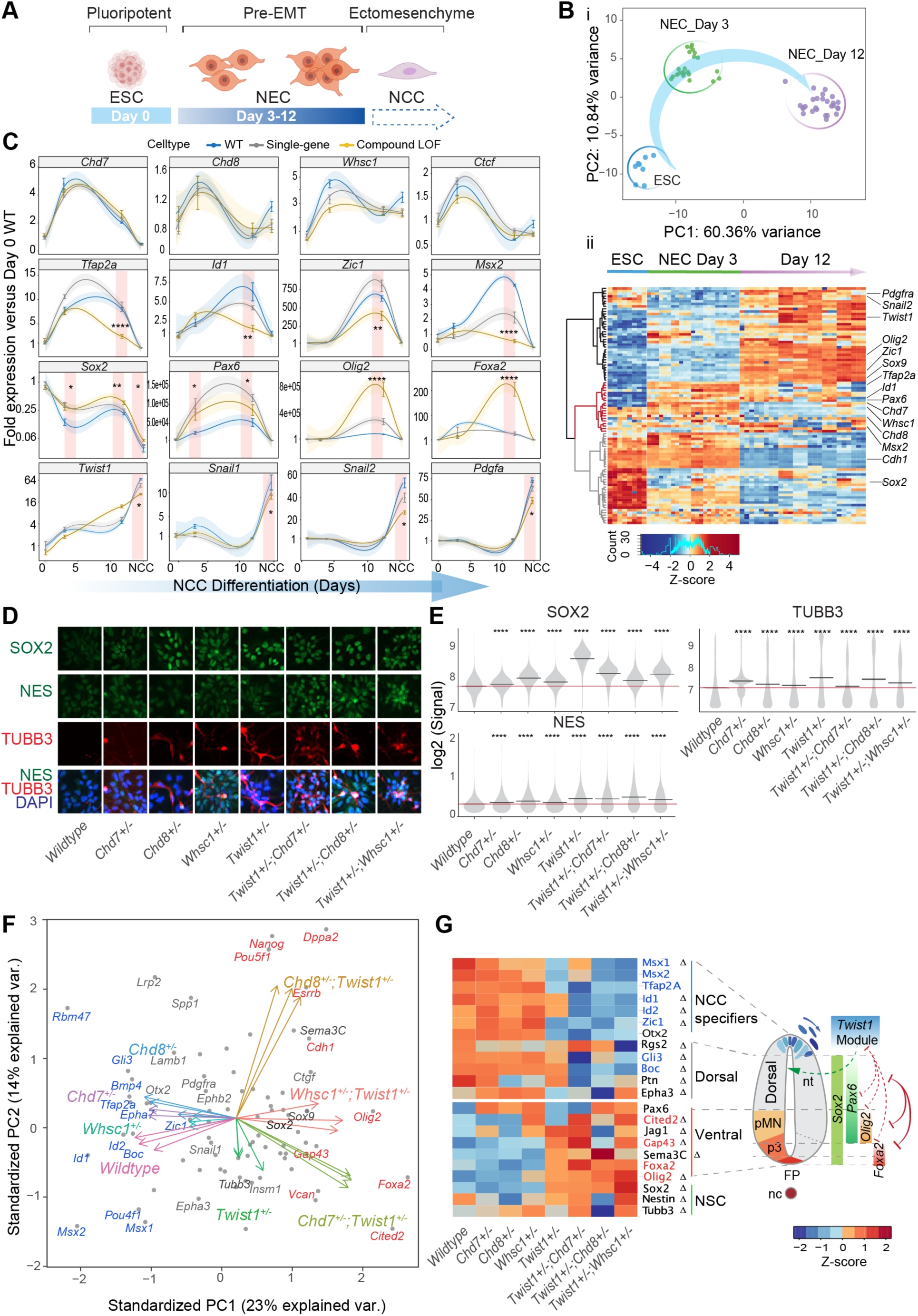
The TWIST1-chromatin regulators predispose NCC propensity and facilitates dorsal-ventral neuroepithelial specification. **A.** Experimental strategy of neural differentiation *in vitro* (Bajpai *et al*., 2010; Varshney *et al*., 2017). **B.** i. Principal component analysis (PCA) of the Fluidigm high-throughput qPCR data for all cell lines collected as ESC, and neuroepithelial cells (NECs) at day 0, day 3 and day 12 of differentiation, respectively. Differentiation trajectory from ESC to NEC is shown for the first two PC axes. ii. Heatmap clustering of normalized gene expressions for all cell lines: n=3 for each genotype analyzed at day 3 and day 12 of neuroepithelium differentiation and n=1 for ESCs. Clusters indicate activated (black), transiently activated (red) and repressed (grey) genes during neural differentiation. Z-score (color-coded) is calculated from log2 transformed normalized expression. **C.** Profiles of expression of representative genes during neural differentiation (day 0 to NCC). Mean expression +/− standard error (SE) are plotted for wildtype, single-gene heterozygous (average of *Twist1^+/−^, Chd7^+/−^, Chd8^+/−^* and *Whsc1^+/−^*) or compound heterozygous (average of *Twist1^+/−^;Chd7^+/−^, Twist1^+/−^;Chd8^+/−^* and *Twist1^+/−^;Whsc1^+/−^*) groups. For NCCs, samples were collected O9-1 cells with siRNA knockdown of single-gene or combinations of *Twist1* and one of the partners. Gene expression were normalized against the mean expression value of 3 housekeeping genes (*Gapdh, Tbp, Actb*), and then the expression of day 0 wildtype ESCs. Shading of trend line represents 90% confidence interval. Red stripes indicate stages when target genes expressions were significantly affected by the double knockdown. P-values were calculated using one-way ANOVA. *P < 0.05, **P < 0.01, ***P < 0.001, **** P < 0.0001. ns, not significant**. D.** Immunofluorescence of SOX2 and selected NSC markers and **E.** quantification of signal intensity +/− SE in single cells of indicated genotypes (X-axis). P-values were generated using one-way ANOVA with Holm-sidak post-test. *P < 0.05, **P < 0.01, ***P < 0.001, **** P < 0.0001. **F.** PCA plot of the NECs (day 12) showing genes with highest PC loadings (blue = top 10 loading, red = bottom 10 loading), and vector of each genotype indicating their weight on the PCs. **G.** Heatmap of genes associated with pattern specification in dorsal-ventral axis of the neuroepithelium: NCC specifiers, dorsal-, ventral-, pan-NSC in mutant versus wildtype NECs. Progenitor identities along the neural tube, the reported master TFs and the co-repression (red solid line) of dorsal and ventral progenitors (pMN, p3 and FP) are illustrated on the right (Briscoe *et al*., 2000; Alaynick *et al*., 2011; Kutejova *et al*., 2016). Repression (red) or promotion (green) of cell fates by TWIST1-module based on the perturbation data are indicated in dashed lines. FP, floor plate; nc, notochord; nt, neural tube. Genes with highest PC loadings were indicated in same colors as in D. Z-scores (color-coded) were calculated from Log2 fold-change against wildtype cells. Changes in gene expressions were significant (by one-way ANOVA). Genes identified as target in at least two ChIP-seq datasets among TWIST1, CHD7 and CHD8 are labelled with Δ.

Gene expression profile of the knockout mutant cells (Figure 7C) were analyzed in conjunction with the gene expression data from NCCs treated with siRNA (representing the “established NCCs”). In the wildtype cells, *Chd7, Chd8, Whsc1* as well as ChIP target *Ctcf*, which encodes a zinc-finger DNA-binding protein that regulates distal promoter-enhancer interactions (Ghirlando and Felsenfeld, 2016), were expressed early in NEC differentiation (Figure 7C, top row). This was followed by an incline in the expression of NCC specifiers (*Tfap2a, Id1, Zic1* and *Msx2;* Figure 7C, second row) and transcription factors (TFs) associated with NSCs (*Sox2, Pax6*, *Olig2*, and *Foxa2;* Figure 7C, third row). A re-activation of chromatin regulators, except for *Chd7*, was observed in the NCCs. The expression of EMT/ ectomesenchyme genes such as *Twist1, Snai1, Snai2* and *Pdgfrα* increased exponentially in established NCCs (Figure 7C, bottom row).

NCC and NSC marker genes responded inversely to the combined perturbation of *Twist1* and chromatin regulators. Compound loss-of-function (LOF) reduced expression of the NCC specifiers and unleashed the expression of NSC TFs in Day-12 NECs (Figure 7C, second and third row). In single-gene heterozygous cells, we observed only modest or no change in the gene expression. *Sox2*, a driver of the NSC lineage and a repressor of NCC formation (Mandalos *et al*., 2014), was repressed concurrently with the increased expression of *Twist1* and the chromatin regulators during neurogenic differentiation and NCC specification (Figure 7C). However, in the compound heterozygous cells, *Sox2* transcript and protein were both up-regulated compared to the wild-type cells, together with NSC markers TUBB3 and NES (Figure 7C-E). Finally, the EMT genes were only affected by the compound knockdown at the established NCC/ ectomesenchyme stage (Figure 7C, bottom row; see also Figure 6D, E).

We focused on effect of gene perturbation on the cell fate bias in late NECs by examining the expression of a broader panel of neural tube patterning genes (Briscoe *et al*., 2000; Alaynick *et al*., 2011; Kutejova *et al*., 2016). The difference between WT and mutant cells in the dataset is primary driven by changes in NCC specifiers and NSC TFs. In the compound mutant NECs, in addition to NCC specifiers (*Tfap2a, Msx1, Msx2, Zic1, Id1* and *Id2)*, expression of dorsal NSC markers were attenuated (*Gli3, Rgs2, Boc* and *Ptn*; Figure 7F, G). Meanwhile, the pan- and ventral-NSC markers *Sox2, Pax6, Olig2, Foxa2* and *Cited2* were ectopically induced (Figure 7F, G: genes in red). ChIP-seq data showed that TWIST1, CHD7 and CHD8 directly bind to the promoters of most of these genes (Figure 4G, S5A, Table S4).

Collectively, the findings implicated that in the plastic NEC progenitor populations, TWIST1-chromatin regulators may help promoting the dorsal fate including the dorsal-most NCC propensity by counteracting SOX2 and other NSC TFs. Loss of function of the module leads to the diversion of NECs from the NCC fate to the neurogenic fate, which may contribute to the deficiency of NCCs and their derivatives observed in the mutant embryos (Figure 3).

## Discussion

### Proteomic screen and network-based inference of NCC epigenetic regulators

Analysis of protein-protein interaction is a powerful approach to identify the connectivity and the functional hierarchy of different genetic determinants associated with an established phenotype (Song and Singh, 2009; Mitra *et al*., 2013; Sahni *et al*., 2015; Cowen *et al*., 2017). We used TWIST1 as an anchor point and the BioID methodology to visualize the protein interactome necessary for NCCs development. Network propagation exploiting similarity network built on prior associations, enabled the extraction of clusters critical for neural crest function and pathology. Using this high throughput analytic pipeline, we were able to identify the core components of the TWIST1-CRM that guides NCC lineage development.

Among the interacting factors were members of the chromatin regulation cluster, which show dynamic component switching between cell types, and may confer tissue-specific activities. The architecture of the modular network reflects the biological organization of chromatin remodeling machinery, which comprises multi-functional subunits with conserved and cell-type-specific components (Meier and Brehm, 2014). Previous network studies reported that disease-causal proteins exist mostly at the center of large clusters and have a high degree of connectivity (Jonsson and Bates, 2006; Ideker and Sharan, 2008). We did not observe an overall correlation between disease probability and the degree of connectivity or centrality for factors in the TWIST1 interactome (Figure S2F). However, the topological characteristics of the chromatin regulatory cluster resembled the features of disease modules and enriched for craniofacial phenotypes. In contrast, the “ribosome biogenesis” module that was also densely inter-connected, was void of relevant phenotypic association (Figure 1C). Network propagation is, therefore, an efficient way to identify and prioritize important clusters while eliminating functionally irrelevant ones.

Based on these results, we selected core TWIST1-CRM epigenetic regulators CHD7, CHD8, and WHSC1, and demonstrated their physical and functional interaction with TWIST1. In the progenitors of the NCCs, these factors displayed overlapping genomic occupancy that correlated with the active chromatin marks in the fate specification genes in neuroepithelium.

### Attribute of TWIST1 interacting partners in NCC development

Combinatorial perturbation of the disease “hot-spots” in TWIST1-CRM impacted adversely on NCC specification and craniofacial morphogenesis in mouse embryos, which phenocopy a spectrum of human congenital malformations associated with NCC deficiencies (Johnson *et al*., 1998; Chun *et al*., 2002; Cai *et al*., 2003; Bosman *et al*., 2005; Bernier *et al*., 2014; Schulz *et al*., 2014; Battaglia *et al*., 2015; Etchevers *et al*., 2019). These observations revealed CHD8 and WHSC1 as putative determinants for NCC development and neurocristopathies. While CHD8 is associated with autism spectrum disorder (Bernier *et al*., 2014; Katayama *et al*., 2016), its function for neural crest development has never been reported. Here, we demonstrated that the loss of *Chd8* affected NCC specification and trigeminal sensory nerve formation *in vivo*, in a *Twist1* dependent manner. We showed that TWIST1 occupancy is a requisite for CHD8 recruitment to common target genes. CHD8 may subsequently initiate chromatin opening and recruit H3-lysine tri-methyltransferases (Zhao *et al*., 2015) such as WHSC1 (Figure S6). We also showed that WHSC1 is required in combination with TWIST1 to promote NCC fate and tissue patterning. Unlike CHD8 and WHSC1, CHD7 has been previously implicated in neurocristopathy (CHARGE syndrome) and the motility of NCCs (Schulz *et al*., 2014; Okuno *et al*., 2017). Our study has corroborated these findings while also showing that CHD7 interacts with TWIST1 to promote NCC specification. In sum, we propose the TWIST1-CRM as a unifying model that connects previously unrelated regulatory factors in different rare diseases and predict their functional dependencies in NCC development (Figure S6). Other epigenetic regulators identified as part of the TWIST1-interactome, such as PBRM1, ZFP62 and MGA, are also part of this module and may act to further fine-tune its activity.

### The phasic activity of TWIST1 and chromatin regulators in the course of NCC differentiation

NCCs are derived from the neuroepithelium in a series of cell fate specification events (Soldatov *et al*., 2019). TWIST1 and the chromatin regulators cooperatively drive the progression along the lineage trajectory at different phases of NCC differentiation. *Twist1* is active at every step as its expression steadily increases during differentiation. The functional interaction with different components of the regulatory module may commence when the expression of *Chd7, Chd8* and *Whsc1* peaks early in the NECs. LOF of the module in NECs leads to enhanced NSC fate bias at the expense of the NCCs, suggesting that the early activation of *Twist1*, *Chd7, Chd8* and *Whsc1* predilect NCC propensity. In the established NCCs, re-activation of *Chd8* and *Whsc1* was associated with the expression of genes associated with NCC identity, EMT, and ectomesenchyme propensity. *Chd7* activity was not coupled with EMT in the NCCs, suggesting that its role may be different from the other two chromatin regulators at this stage. The switch of TWIST1 module activity was reflected by activation of different groups of target genes, suggesting that phase-specific deployment of the regulatory module is critical for navigating the cells along the lineage trajectory of NCC development.

### The competition between TWIST1 module and SOX2 in fate decision

The segregation of NCC and NSC lineages is the first event of NCC differentiation. Our results show that the lineage allocation may be accomplished by the mutual opposition between core members of the TWIST1-CRM and NSC TFs such as SOX2. *Sox2* expression is continuously repressed in the NCC lineage (Wakamatsu *et al*., 2004; Cimadamore *et al*., 2011; Soldatov *et al*., 2019), likely through direct binding and inhibition by TWIST1-CHD8 at *Sox2* promotor. In *Twist1^+/−^;Chd8^+/−^* mutant embryos, the aberrant upregulation of *Sox2* correlated with deficiency of NCC derivatives and the expanded neuroepithelium of the embryonic brain. In a similar context, *Sox2* overexpression in chicken neuroepithelium blocks the production of TFAP2α-positive NCC (resulting in the loss of cranial nerve ganglia), and circumvented the expression of EMT genes and NCC ventral migration (Wakamatsu *et al*., 2004; Remboutsika *et al*., 2011). On the contrary, conditional knockout of *Sox2* results in ectopic formation and migration of NCCs and thinning of the neuroepithelium (Mandalos *et al*., 2014).

### The TWIST1-chromatin regulator induces NCC specification concurrently with dorsoventral polarization of the NSCs

The partitioning of the neural tube into subdomains along the dorsoventral axis is accomplished by cross-repression of the domain-specific transcriptional activity (Briscoe *et al*., 2000; Kutejova *et al*., 2016). Our findings suggest a secondary contribution of the TWIST1-chromatin regulators to this process. Our data suggest that when NCC and NSC fate programs are activated in the neural progenitors, TWIST1-chromatin regulators repress both pan- and ventral-NSC TFs (*Sox2*, *Olig2, Foxa2, Pax6*) and their effectors while concurrently promoting the dorsal neuroepithelial fate necessary for priming the cells for NCC specification. Reciprocally, NSC transcription factors, including SOX2, PAX6, and OLIG2 (Hikichi *et al*., 2013; Mistri *et al*., 2015; Kutejova *et al*., 2016) may repress the TWIST1-chromatin regulators and compete with them at the promotors of NCC specifiers to enhance NSC fate (Figure S5B, Figure S6). Notably, SOX2 and PAX6 are expected to co-bind and activate genes promoting the ventral fate (Zhang *et al*., 2019). These results indicate that the NCC lineage is established concurrently with neurogenic lineages and that it is abided by the same patterning rules. The TWIST1-CRM members may therefore also be part of the molecular machinery necessary for dorsoventral partitioning of the neuroepithelium (Briscoe *et al*., 2000; Kutejova *et al*., 2016)..

In conclusion, by implementing an analytic pipeline to decipher the TWIST1 interactome, we have a glimpse of the global molecular hierarchy of NCC development. We have characterized the cooperative function of core components of TWIST1-CRM including the TWIST1 and chromatin regulators CHD7, CHD8 and WHSC1. We demonstrated that this module is a dynamic nexus to drive molecular mechanisms for orchestrating NCC lineage progression and repressing NSC fate, facilitating dorsoventral tissue patterning and enabling the acquisition of ectomesenchyme propensity. The TWIST1-chromatin regulators and the NSC regulators coordinate the cross-talk between the neural crest-derived tissues and the neural progenitors of the CNS, both of which are often affected concurrently in a range of human congenital diseases.

## Materials and Methods

### Cell culture and BioID Protein proximity-labeling

O9-1 cells (passage 20-22, Millipore cat. #SCC049) were maintained in O9-1 medium: high glucose DMEM (Gibco), 12.5 % (v/v) heat-inactivated FBS (Fisher Biotec), 10 mM β-mercaptoethanol, 1X non-essential amino acids (100X, Thermo Fisher Scientific), 1 % (v/v) nucleosides (100X, Merck) and 10 mil U/mL ESGRO® mouse leukaemia inhibitory factor (Merck) and 25 ng/mL FGF-2 (Millipore, Cat. #GF003). For each replicate experiment, 1.5 x10^6^ cells per flask were seeded onto 4*T75 flasks 24 hrs before transfection. The next day *PcDNA 3.1/ Twist1-BirA*-HA* plasmid or *PcDNA 3.1/ GFP-BirA*-HA* plasmid was transfected into cells using Lipofectamine® 3000 (Life Technologies) according to the manufacturer’s instructions. Biotin (Thermo Scientific, cat. #B20656) was applied to the medium at 50 nM. Cells were harvested 16 hrs post-transfection, followed by snap-freeze liquid nitrogen storage or resuspension in lysis buffer. All steps were carried out at 4°C unless indicated otherwise. Cells were sonicated on the Bioruptor Plus (Diagenode), 30s on/off for five cycles at high power. An equal volume of cold 50 mM Tris-HCl, pH 7.4, was added to each tube, followed by two 30s on/off cycles of sonication. Lysates were centrifuged for 15 mins at 14000 rpm. Protein concentrations were determined by Direct Detect® Infrared Spectrometer (Merck).

Cleared lysate with equal protein concentration for each treatment was incubated with pre-blocked streptavidin Dynabeads® (MyOne Streptavidin C1, Invitrogen™, cat. #65002) for 4 hrs. Beads were collected and washed sequentially in Wash Buffer 1-3 with 8 mins rotation each, followed by quick washes with cold 1 mL 50 mM Tris·HCl, pH 7.4, and 500 μL triethylammonium bicarbonate (75 mM). Beads were then collected by spinning (5 min at 2,000 × *g)* and processed for mass spectrometry analysis.

*<u>Lysis Buffer:</u>* 500mM NaCl, 50 mM Tris-HCl, pH 7.5, 0.2% SDS 0.5% Triton. Add 1x Complete protease inhibitor (Roche), 1 mM DTT fresh

*<u>Wash Buffer 1</u>*: 2 % SDS,

*<u>Wash Buffer 2</u>*: 0.1 % sodium deoxycholate, 1% Triton, 1mM EDTA, 1 mL, 500 mM NaCl, 50 mM HEPES-KOH, pH 7.5

*<u>Wash Buffer 3</u>*: 0.1 % sodium deoxycholate, 0.5% NP40 (Igepal), 1 mM EDTA, 250 mM NaCl, 10 mM Tris-HCl, pH 7.5

### Liquid chromatography with tandem mass spectrometry

Tryptic digestion of bead-bound protein was performed in 5% w/w trypsin (Promega, cat. #V5280), 50mM triethylammonium bicarbonate buffer at 37 °C overnight. The supernatant was collected and acidified with trifluoroacetic acid (TFA, final concentration 0.5% v/v). Proteolytic peptides were desalted using Oligo R3 reversed phase resin (Thermo Fisher Scientific) in stage tips made in-house (Rappsilber *et al*., 2007). Peptides were fractioned by hydrophilic interaction liquid chromatography using an UltiMate 3000 HPLC (Thermo Fisher Scientific) and a TSKgel Amide-80 HILIC 1 mm × 250 mm column. Peptides were eluted in a gradient from 100% mobile phase B (90% acetonitrile, 0.1% TFA, 9.9% water) to 60% mobile phase A (0.1% TFA, 99.9% water) for 35 min at 50 µL/min and fractions collected in a 96-well plate, followed by vacuum centrifugation to dryness. Dried peptide pools were reconstituted in 0.1% formic acid in the water, and 1/10th of samples were analyzed by LC-MS/MS.

Mass spectrometry was performed using an LTQ Velos-Orbitrap MS (Thermo Fisher Scientific) coupled with an UltiMate RSLCnano-LC system (Thermo Fisher Scientific). A volume of 5 µL was loaded onto a 5 mm C18 trap column (Acclaim PepMap 100, 5 µm particles, 300 µm inside diameter, Thermo Fisher Scientific) at 20 µL/ min for 2.5 min in 99% phase A (0.1% formic acid in water) and 1% phase B (0.1% formic acid, 9.99% water and 90% acetonitrile). The peptides were eluted through a 75 µm inside diameter column with integrated laser-pulled spray tip packed to a length of 20 cm with Reprosil 120 Pur-C18 AQ 3 µm particles (Dr. Maisch). The gradient was from 7% phase B to 30% phase B in 46.5 min, to 45% phase B in 5 min, and to 99% phase B in 2 min. The mass spectrometer was used to apply 2.3 kV to the spray tip via a pre-column liquid junction. During each cycle of data-dependent MS detection, the ten most intense ions within m/z 300-1,500 above 5000 counts in a 120,000 resolution orbitrap MS scan were selected for fragmentation and detection in an ion trap MS/MS scan. Other MS settings were: MS target was 1,000,000 counts for a maximum of 500 ms; MS/MS target was 50,000 counts for a maximum of 300 ms; isolation width, 2.0 units; normalized collision energy, 35; activation time 10 ms; charge state 1 was rejected; mono-isotopic precursor selection was enabled; dynamic exclusion was for 10 s.

### Proteomic data analysis

#### Pre-processing of raw mass spectrometry data

Raw MS data files were processed using Proteome Discoverer v.1.3 (Thermo Fisher Scientific). Processed files were searched against the UniProt mouse database (downloaded Nov 2016) using the Mascot search engine version 2.3.0. Searches were done with tryptic specificity allowing up to two missed cleavages and tolerance on mass measurement of 10 ppm in MS mode and 0.3 Da for MS/MS ions. Variable modifications allowed were acetyl (Protein N-terminus), oxidized methionine, glutamine to pyro-glutamic acid, and deamidation of asparagine and glutamine residues. Carbamidomethyl of cysteines was a fixed modification. Using a reversed decoy database, a false discovery rate (FDR) threshold of 1% was used. The lists of protein groups were filtered for first hits.

Processing and analysis of raw peptide-spectrum match (PSM) values were performed in R following the published protocol (Waardenberg, 2017). Data were normalized by the sum of PSM for each sample (Figure S2B), based on the assumption that the same amount of starting materials was loaded onto the mass spectrometer for the test and control samples. A PSM value of 0 was assigned to missing values for peptide absent from the sample or below detection level (Sharma *et al*., 2009). Data points filtered by the quality criterion that peptides had to be present in at least two replicate experiments with a PSM value above 2. The normalized and filtered dataset was fitted under the negative binomial generalized linear model and subjected to the likelihood ratio test for TWIST1 vs. control interactions, using the msmsTest and EdgeR packages (Gregori *et al*., 2019) (Robinson *et al*., 2010). Three biological replicates each from O9-1 and 3T3 cells were analyzed. One set of C3H10T1/2 cell line was analyzed. A sample dispersion estimate was applied to all datasets. Stringent TWIST1-specific interactions in the three cell lines were determined based on a threshold of multi-test adjusted p-values (adjp) < 0.05 and fold-change > 3.

#### Network propagation for functional identification and novel disease gene annotation

Prior knowledge of mouse protein functional associations, weighted based on known protein-protein interaction (PPI), co-expression, evolutionary conservation, and test mining results, were retrieved by the Search Tool for the Retrieval of Interacting Genes (STRING) (Szklarczyk *et al*., 2015). Intermediate confidence (combined score) of > 0.4 was used as the cut-off for interactions. The inferred network was imported into Cytoscape for visualization (Shannon *et al*., 2003). We used MCL algorism (Enright *et al*., 2002), which emulates random walks between TWIST1 interacting proteins to detect clusters in the network, using the STRING association matrix as the probability flow matrix. Gene Ontology and transcriptional binding site enrichment analysis for proteins were obtained from the ToppGene database (Chen *et al*., 2009), with a false-discovery rate < 0.05. The enriched functional term of known nodes was used to annotate network neighbors within the cluster with unclear roles.

Heat diffusion was performed on the network, using twenty-two genes associated with human and mouse facial malformation (HP:0001999, MP:0000428) as seeds. A diffusion score of 1 was assigned to the seeds, and these scores were allowed to propagate to network neighbors, and heat stored in nodes after set time = 0.25 was calculated. NetworkAnalyzer (Assenov *et al*., 2008), which is a feature of Cytoscape, was used to calculate nodes’ Degree (number of edges), Average Shortest Path (connecting nodes), and Closeness Centrality (a measure of how fast information spreads to other nodes).

### Co-immunoprecipitation

#### Protein Immunoprecipitation

For the analysis of protein localization, transfection was performed using Lipofectamine® 3000 (Life Tech) according to manufacturer instructions with the following combinations of plasmids: pCMV-*Twist1-FLAG* plus one of (*pCMV-gfp-HA*, *pCMV-Tcf3-HA, pCMV-Prrx1-HA, pCMV-Prrx2-HA, pCMV-Chd7-HA, pCMV-Chd8-HA, pCMV-Dvl1-HA, pCMV-Smarce1-HA, pCMV-Tfe3-HA, pCMV-Whsc1-HA, pCMV-Hmg20a-HA).* The cell pellet was lysed and centrifuged at 14,000 × *g* for 15 min. Cleared lysate was incubated with *α*-TWIST1/ *α*-FLAG antibody (1 μg/ mL) at 4°C for 2 hrs with rotation. Protein-G agarose beads (Roche) were then added, and the sample rotated for 30 min at RT °C. Beads were washed in ice-cold wash buffer six times and transferred to new before elution in 2x LDS loading buffer at 70°C for 10 mins. Half the eluate was loaded on SDS-PAGE with the “input” controls for western blot analysis.

#### Western Blotting

Protein was extracted using RIPA buffer lysis (1× PBS, 1.5% Triton X-100, 1% IGEPAL, 0.5% Sodium Deoxycholate, 0.1% SDS, 1 mM DTT, 1x Complete protease inhibitor [Roche]) for 30 minutes at 4°C under rotation. The lysate was cleared by centrifugation at 15000 g, and protein concentration was determined using the Direct Detect spectrometer (Millipore). 20 µg of protein per sample was denatured at 70°C for 10 mins in 1× SDS Loading Dye (100 mM Tris pH 6.8, 10% (w/v) SDS, 50% (w/v) Glycerol, 25%(v/v) 2-Mercaptoethanol, Bromophenol blue) and loaded on a NuPage 4-12% Bis-Tris Gel (Life Technologies, Cat. #NP0322BOX). Electrophoresis and membrane transfer was performed using the Novex™ (Invitrogen) system following manufacturer instructions.

Primary antibodies used were mouse monoclonal *α*-TWIST1 (1:1000, Abcam, Cat. #ab50887), mouse monoclonal [29D1] *α*-WHSC1/NSD2 (1:5000, Abcam, Cat. #ab75359), rabbit polyclonal *α*-CHD7 (1:5000, Abcam, Cat. #ab117522), rabbit polyclonal *α*-CHD8 (1:10000, Abcam, Cat. #ab114126), mouse *α*-α-tubulin (1:1000, Sigma, Cat. #T6199), rabbit *α*-HA (1:1000, Abcam, Cat. #ab9110) and mouse *α*-FLAG M2 (Sigma, Cat. #F1804). Secondary antibodies used were HRP-conjugated donkey *α*-Rabbit IgG (1:8000, Jackson Immunoresearch, Cat. #711-035-152) and HRP-conjugated donkey *α*-Mouse IgG (1:8000, Jackson Immunoresearch, Cat. #711-035-150).

### GST Pull-down

#### Production and purification of recombinant proteins

Prokaryotic expression plasmids pGEX2T with the following inserts GST-*Twist1*, GST-*N’Twist1*, GST-*C’Twist1*, GST-*Twist1bhlh*, GST-*Twist1TA, or* GST were transfected in BL21 (DE3) *Escherichia coli* bacteria (Bioline). Bacterial starter culture was made by inoculation of 4 mL Luria broth with 10 μg/mL ampicillin, and grown 37°C, 200 rpm overnight. Starter culture was used to inoculate 200 mL Luria broth media with 10 μg/mL ampicillin and grown at 37°C, 200 rpm until the optical density measured OD600 was around 0.5-1.0. The culture was cooled down to 25°C for 30 min before Isopropyl β-D-1-thiogalactopyranoside (IPTG) was added to the media at a final concentration of 1 mM. Bacteria were collected by centrifugation 4 hrs later at 8000 rpm for 10 mins at 4°C.

Bacteria were resuspended in 5 % volume of lysis buffer (10 mM Tris-Cl, pH 8.0; 300 mM NaCl; 1 mM EDTA, 300 mM NaCl, 10 mM Tris.HCl [pH 8.0], 1 mM EDTA, 1x Complete protease inhibitor [Roche], 1 mM PMSF, 100ng/mL leupeptin, 5mM DTT) and nucleus were released by 3 rounds of freeze/ thaw cycles between liquid nitrogen and cold water. The sample was sonicated for 15 s × 2 (consistent; intensity 2), with 3 min rest on ice between cycles. Triton X-100 was added to a final concentration of 1%. The lysate was rotated for 30 min at 4°C and centrifuged at 14000 rpm for 15 min at 4°C.

The supernatant was collected and rotated with 800 μL of 50% Glutathione Sepharose 4B slurry (GE, cat. # 17-0756-01) for 1 h, at 4°C. Beads were then loaded on MicroSpin columns (GE cat. #27-3565-01). Column was washed three times with wash buffer (PBS 2X, Triton X-100 0.1%, imidazole 50 mM, NaCl 500 mM, DTT 1 mM, 1x Complete protease inhibitor [Roche]) before storage in 50% glycerol (0.01 % Triton). Quantity and purity of the recombinant protein on beads were assessed by SDS polyacrylamide gel electrophoresis (SDS-PAGE, NuPAGE 4-12 % bisacrylamide gel, Novex) followed by Coomassie staining or western blot analysis with anti-TWIST1 (1:1000), anti-GST (1:1000) antibody. Aliquots were kept at −20°C for up to 6 months.

#### GST pulldown

Cell pellet (5 × 10^6^) expressing HA-tagged TWIST1 interaction candidates were thawed in 300 μL hypotonic lysis buffer (HEPES 20 mM, MgCl2 1 mM, Glycerol 10%, Triton 0.5 %. DTT 1 mM, 1x Complete protease inhibitor [Roche], Benzo nuclease 0.5 μl/ ml) and incubated at room temperature for 15 mins (for nuclease activity). An equal volume of hypertonic lysis buffer (HEPES 20 mM, NaCl2 500 mM, MgCl2 1 mM, Glycerol 10%, DTT 1 mM, 1x Complete protease inhibitor [Roche]) was then added to the lysate. Cells are further broken down by passaging through gauge 25 needles for 10 strokes and rotated at 4°C for 30 min. After centrifugation at 12,000 × g, 10 min, 200 μL lysate was incubated with 10 μL bead slurry (or the same amount of GST fusion protein for each construct decided by above Coomassie staining). Bait protein capture was done at 4°C for 4 hrs with rotation.

Beads were collected by spin at 2 min at 800 × g, 4°C, and most of the supernatant was carefully removed without disturbing the bead bed. Beads were resuspended in 250 μL ice-cold wash buffer, rotated for 10 mins at 4°C and transferred to MicroSpin columns that were equilibrated with wash buffer beforehand. Wash buffer was removed from the column by spin 30 sec at 100 × g, 4°C. Beads were washed for 4 more times quickly with ice-cold wash buffer before eluting proteins in 2X LDS loading buffer 30 μL at 70°C, 10 min, and characterized by western blotting.

### Generation of mutant ESC by CRISPR-Cas9 editing

CRISPR-Cas9-edited mESCs were generated as described previously (Sibbritt *et al*., 2019). Briefly, 1-2 gRNAs for target genes were ligated into pSpCas9(BB)-2A-GFP (PX458, addgene plasmid #48138*, a gift from Feng Zhang)*. Three µg of pX458 containing the gRNA was electroporated into 1×10^6^ A2loxCre ESCs or A2loxCre Twist1+/− cells (clone T2-3, generated by the Vector & Genome Engineering Facility at the Children’s Medical Research Institute) using the Neon® Transfection System (Thermo Fisher Scientific). Electroporated cells were plated as single cells onto pre-seeded lawns of mouse embryonic fibroblasts (MEF), and GFP expressing clones grown from single cells were selected under the fluorescent microscope. In total, 30-40 clones were picked for each electroporation. For mutant ESC genotyping, clones were expanded and grown on a gelatin-coated plate for three passages, to remove residue MEFs contamination.

For genotyping, genomic lysate of ESCs was used as input for PCR reaction that amplified region surrounding the mutation site (+/− 200-500 bp flanking each side of the mutation). The PCR product was gel purified and sub-cloned into the pGEM®-T Easy Vector System (Promega) as per manufacturer’s protocol. At least ten plasmids from each cell line were sequenced to ascertain monoallelic frameshift mutation and exclude biallelic mutations.

### Generation of mouse chimeras from ESCs

ARC/s and *DsRed.T3* mice were purchased from the Australian Animal Resources Centre and maintained as homozygous breeding pairs. ESC clones with monoallelic frameshift mutations and the parental *A2LoxCre* ESC line were used to generate chimeras. Embryo injections were performed as previously described (Sibbritt *et al*., 2019). Briefly, 8-10 ESCs were injected per eight-cell *DsRed.T3* embryo (harvested at 2.5 dpc from super-ovulated *ARC/s* females crossed to *DsRed.T3* stud males) and incubated overnight. Ten to twelve injected blastocysts were transferred to each E2.5 pseudo-pregnant ARC/s female recipient. E9.5 and E11.5 embryos were collected 6 and 8 days after transfer to pseudo-pregnant mice. Embryos showing red fluorescent signal indicating no or low ESC contribution were excluded from the phenotypic analysis. Animal experimentations were performed in compliance with animal ethics and welfare guidelines stipulated by the Children’s Medical Research Institute/Children’s Hospital at Westmead Animal Ethics Committee.

### Whole-mount fluorescent immunostaining of mouse embryos

Whole-mount fluorescent immunostaining of mouse embryos was performed by following the procedure of (Adameyko *et al*., 2012) with minor modifications. Embryos were fixed for 6 hours in 4% paraformaldehyde (PFA) and dehydrated through a methanol gradient (25%, 50%, 75%, 100%). After 24 hours of incubation in 100% methanol at 4°C, embryos were transferred into bleaching solution (1 part of 30% hydrogen peroxide to 2 parts of 100% methanol) for another 24 hours (4°C). Embryos were then washed with 100% methanol (10 minutes x3 at room temperature), post-fixed with Dent’s Fixative (dimethyl sulfoxide: methanol = 1:4) overnight at 4°C.

Embryos were blocked for 1 hour on ice in blocking solution (0.2 % BSA, 20% DMSO in PBS) with 0.4% Triton. Primary antibodies mouse 2H3 (for neurofilament 1:1000) and rabbit *α*-TFAP2A (1:1000) or were diluted in blocking solution and incubated for four days at room temperature, and secondary antibodies (Goat *α*-Rabbit Alexa Fluor 633; Goat *α*-Mouse Alexa Fluor 488 and DAPI, Thermo Fisher Scientific) were incubated overnight in blocking solution at room temperature. Additional information of the antibodies used are listed in Table S6. Embryos were cleared using BABB (1part benzyl alcohol: 2 parts benzyl benzoate), after dehydration in methanol, and imaged using a Carl Zeiss Cell Observer SD spinning disc microscope. Confocal stacks through the embryo were acquired and then collapsed. Confocal stacks were produced containing ∼150 optical slices. Bitplane IMARIS software was used for 3D visualization and analysis of confocal stacks. Optical sections of the 3D embryo were recorded using ortho/oblique functions in IMARIS software. The surface rendering wizard tool was used to quantify SOX2 expression in the ventricular zone by measuring the immunofluorescence intensity on three separate z-plane sections per volume of the region of each embryo. The data were presented graphically as the ratio of intensity/ volume.

### Generation of TWIST1 inducible expression ESC line

ESC lines generated are listed in Table S6. A2loxCre Mouse ESCs (Mazzoni *et al*., 2011) was a gift from Kyba Lab (Lillehei Heart Institute, Minnesota, USA). A2loxCre with Twist1 bi-allelic knockout background was generated by CRISPR-Cas9, as described below. The inducible *Twist1* ESC line was generated using the inducible cassette exchange method described previously (Iacovino *et al*., 2014). The TWIST1 coding sequence was then cloned from the mouse embryo cDNA library into the p2lox plasmid downstream of the Flag tag (Iacovino *et al*., 2014). The plasmid was transfected into A2loxCre (*Twist1 ^−/−^*) treated with 1 μg/mL doxycycline for 24 hrs. The selection was performed in 300 μg/mL of G418 (Gibco) antibiotic for one week. Colonies were then picked and tested for TWIST1 expression following doxycycline treatment.

### NEC differentiation of the ESCs

ESC lines generated in this study were differentiated into neural epithelial cells (NECs) following established protocols (Bajpai *et al*., 2010; Varshney *et al*., 2017) with minor modifications. ESCs were expanded in 2i/LIF media (Ying *et al*., 2008) for 2-3 passages. Neurogenic differentiation was initiated by plating ESC in AggreWells (1×10^6^ per well) using feeder independent mESC. Colonies were then lifted from AggreWells and grown in suspension in Neurogenic Differentiation Media supplemented with 15% FBS with gentle shaking for 3 days. Cell colonies were transferred to gelatin-coated tissue culture plates and cultured for 24 h at 37 °C under 5% CO_2_.

Cells were selected in insulin-transferrin-selenium (ITS)-Fibronectin media for 6-8 days at 37 °C and 5% CO_2_, with a change of media every other day. Accutase^TM^ (Stemcell Technologies) was used to dissociate cells from the plate, allowing the removal of cell clumps. NECs were collected by centrifugation and plated on Poly-L-ornithine (50 μg/mL, Sigma-Aldrich) and Laminin (1 μg/mL, Novus Biological) coated dishes. For expansion of the cell line, cells were cultured in Neural Expansion Media (1.5 mg/mL Glucose, 73 μg/mL L-glutamine, 1x N2 media supplement [R & D systems] in Knockout DMEM/F12 [Invitrogen], 10 ng/mL FGF-2 and 1 μg/mL Laminin [Novus Biologicals]). During this period, cells were lifted using Accutase^TM^ and cell rosette clusters were let settle and were removed for two passages to enrich for pre-EMT NCC populations.

### Chromatin immunoprecipitation Sequencing (ChIP-seq)

ESC with genotype *Twist1*^−/−^; *Flag-Twist1* O/E and *Twist1*^−/−^ were differentiated into NEC for 3 days following established protocol (Varshney *et al*., 2017) and were collected in ice-cold DPBS. Following a cell count, approximately 2 × 10^7^ cells were allocated per cell line per ChIP. ChIP-seq assays were performed as previously described (Bildsoe *et al*., 2016). In brief, chromatin was crosslinked and sonicated on the Bioruptor Plus (Diagenode) using the following program: 30 seconds on/off for 40 minutes on High power. The supernatant was incubated with *α*-TWIST1 (Abcam, at. #ab50887) antibody conjugated Dynabeads overnight at 4 °C. The protein-chromatin crosslinking is reversed by incubation at 65 °C for 6 h. The DNA is purified using RNase A and proteinase K treatments, extracted using phenol-chloroform-isoamyl alcohol (25:24:1, v/v) and precipitated using glycogen and sodium acetate. The precipitated or input chromatin DNA was purified and converted to barcoded libraries using the TruSeq ChIP Sample Prep Kit (Illumina). Then 101 bp paired-end sequencing was performed on the HiSeq 4000 (Illumina).

### ChIP-sequencing data analysis

ChIP-seq quality control results and analysis can be found in Figure S4. Adaptors from raw sequencing data were removed using Trimmomatic (Bolger *et al*., 2014) and aligned to the *mm10* mouse genome (GENCODE GRCm38.p5; (Frankish *et al*., 2019) using BWA aligner (Li and Durbin, 2009), and duplicates/unpaired sequences were removed using the picardtools (http://broadinstitute.github.io/picard/). MACS2 package (Zhang *et al*., 2008) was used for ChIP-seq peak calling for both *Twist1^−/−^; Flag-Twist1 O/E* and *Twist1^−/−^* IP samples against genomic input. IDR analysis was performed using the P-value as the ranking measure, with an IDR cut-off of 0.05. Peak coordinates from the two replicates were merged, using the most extreme start and end positions. The raw and processed data were deposited into the NCBI GEO database and can be accessed with the accession number GSE130251.

### ChIP-seq integrative analysis

Public ChIP-seq datasets for CHD7, CHD8 and histone modifications in NECs were selected based on the quality analysis from the Cistrome Data Browser (http://cistrome.org/db/#/) and ENCODE guideline (Encode, 2012; Mei *et al*., 2017). Datasets imported for analysis are listed in Table S6. To facilitate comparison with datasets generated from human samples, TWIST1 ChIP sequences were aligned to the hg38 human genome by BWA. ChIP peak coordinates from this study were statistically compared using fisher’s exact test (cut-off: P-value < 0.05, odds ration >10) and visualized using Jaccard similarity score. Analysis were performed with BEDTools (Quinlan and Hall, 2010). ChIP-seq peaks for TWIST1, CHD7 and CHD8 were extended to uniform 1 kb regions, and regions bound by single factors or co-occupied by 2 or 3 factors were identified. The Genomic Regions Enrichment of Annotations Tool (GREAT) was used to assigns biological functions to genomic regions by analyzing the annotations of the nearby genes (Mclean *et al*., 2010). Significance by both binomial and hypergeometric test (P<0.05) were used as cut-off. Genes with TSS +/− 5 kb of the peaks were annotated using ChIPpeakAnno package in R. List of target genes was compared between each CHD7, CHD8, and TWIST1. Bam files for each experiment were converted to bigwig files for ChIP-seq density profile, chromosome footprint, and IGV track visual analysis.

### O9-1 siRNA treatment and scratch assay

Scratch Assays were performed on O9-1 cells following transient siRNA lipofectamine transfections. O9-1 cells were seeded at a density of 0.5×10^5^ cells per well on Matrigel-coated 24-well plates on the day of transfection. 20 pmol of siRNA for candidate gene (*Chd7, Chd8* or *Whsc1*) and 20 pmol siRNA for *Twist1* or control was applied per well (24-well-plate), plus 3 μL lipofectamine RNAiMAX reagent (Thermo Fisher Scientific, cat. #13778075), following manufacturer protocol. Knockdown efficiency was assessed by qRT-PCR (Figure S5).

48 hours after transfection, a scratch was made in the confluent cell monolayer. Live images were taken with the Cell Observer Widefield microscope (ZEISS international) under standard cell culture conditions (37°C, 5% CO2). Bright-field images were captured at set tile regions every 15 mins over a 10 hrs period. The total migration area from the start of imaging to when the first cell line closed the gap was quantified by Fiji software (Schindelin *et al*., 2012).

### cDNA synthesis, pre-amplification, and Fluidigm high-throughput RT-qPCR analysis

cDNA synthesis, from 1 µg total RNA from each sample, was performed using the RT2 Microfluidics qPCR Reagent System (Qiagen, Cat. # 330431). cDNAs were pre-amplified using the primer Mix for reporter gene sets (Table S5). High-throughput gene expression analysis (BioMarkTM HD System, Fluidigm) was then performed using the above primer set.

Raw data were extracted using the Fluidigm Real-Time PCR Analysis Software, and subsequent analysis was performed in R-studio. Ct values flagged as undetermined or higher than the threshold (Ct > 24) were assigned as missing values. Samples with a measurement for only one housekeeping gene or samples with measurements for < 30 genes were excluded from further analysis. Genes missing values for more than 30 samples were also excluded from further analysis. Data were normalized using expressions of the average of 3 housekeeping genes (*Gapdh, Tbp, Actb*). Regularized-log transformation of the count matrix was then performed, and the PCA loading gene was generated using functions in the DEseq2 package. Differential gene expression analysis was performed using one-way ANOVA.

## Acknowledgment

Our work was supported by the National Health and Medical Research Council (NHMRC) of Australia (Grant ID 1066832), the Australian Research Council (Grant DP 1094008) and Mr. James Fairfax (Bridgestar Pty Ltd). Imaging analysis was performed at the ACRF Telomere Analysis Centre and proteomics analysis was performed at the Biomedical Proteomics Facility, both supported by the Australian Cancer Research Foundation. XCF was supported by the University of Sydney International Postgraduate Research Scholarship, the Australian Postgraduate Award and the CMRI Scholarship; KEK was supported by The Danish Council for Independent Research and FP7 Marie Curie Actions – COFUND (DFF – 1325-00154) and the Carlsberg Foundation (CF15-1056 and CF16-0066); PO is the CMRI Norman Gregg Research Fellow; MEG was supported by NHMRC (grant ID 1079160); NF was supported by a University of Sydney Post-Doctoral Fellowship and the CMRI Norman Gregg Research Fellowship and PPLT is an NHMRC Senior Principal Research Fellow (Grant ID 1003100, 1110751).

## Author contribution

X.F., N.F. and P.P.L.T. designed the project; X.F., P.M., and J.Q.J.S. conducted the experiments; M.G. K.E.K. provided technical assistance with proteomics experiment and analysis, P.O. and J.S. assisted the transcriptome and imaging experiment; X.F. performed the bioinformatics analysis; X.F., N.F. and P.P.L.T. wrote the manuscript. All authors edited the manuscript.

## Supplemental Figures

**Figure S1.**
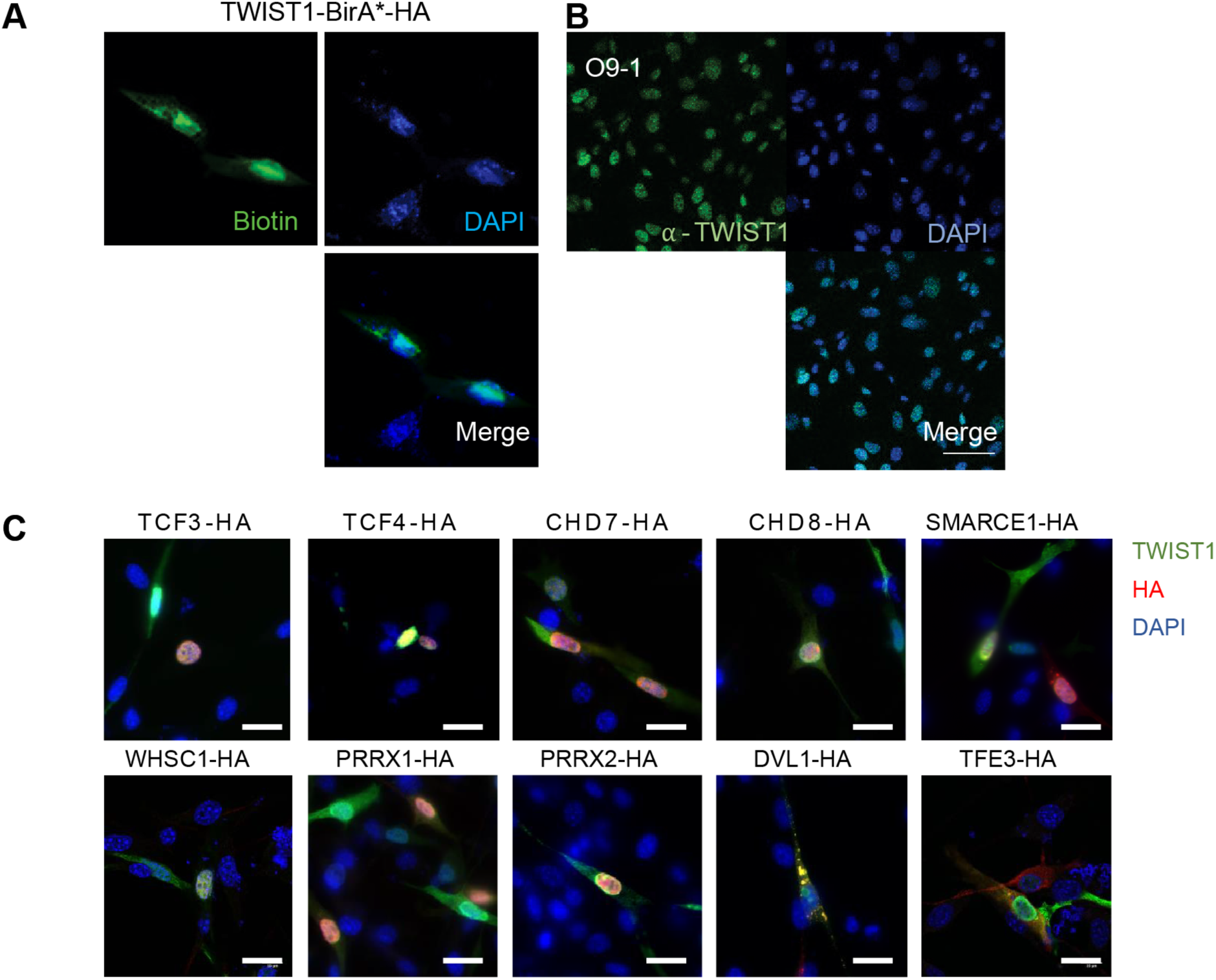
Nuclear localization of TWIST1-BirA* biotinylated proteins recapitulates that of the endogenous TWIST1. **A.** Immunofluorescence analysis revealed co-localization of TWIST1-BirA* (HA tagged) and biotinylated proteins (labelled with streptavidin-GFP) in the nucleus in NCCs. Bar = 20 μm. **B.** Expression and localization of endogenous TWIST1 in untransfected cells stained by *α*-TWIST1. Bar = 50 μm. **C.** Immunofluorescence detection of proteins in cells co-expressing FLAG-TWIST1 (α-TWIST1) and HA-tagged proteins (α-HA). Nuclei were stained by DAPI.

**Figure S2.**
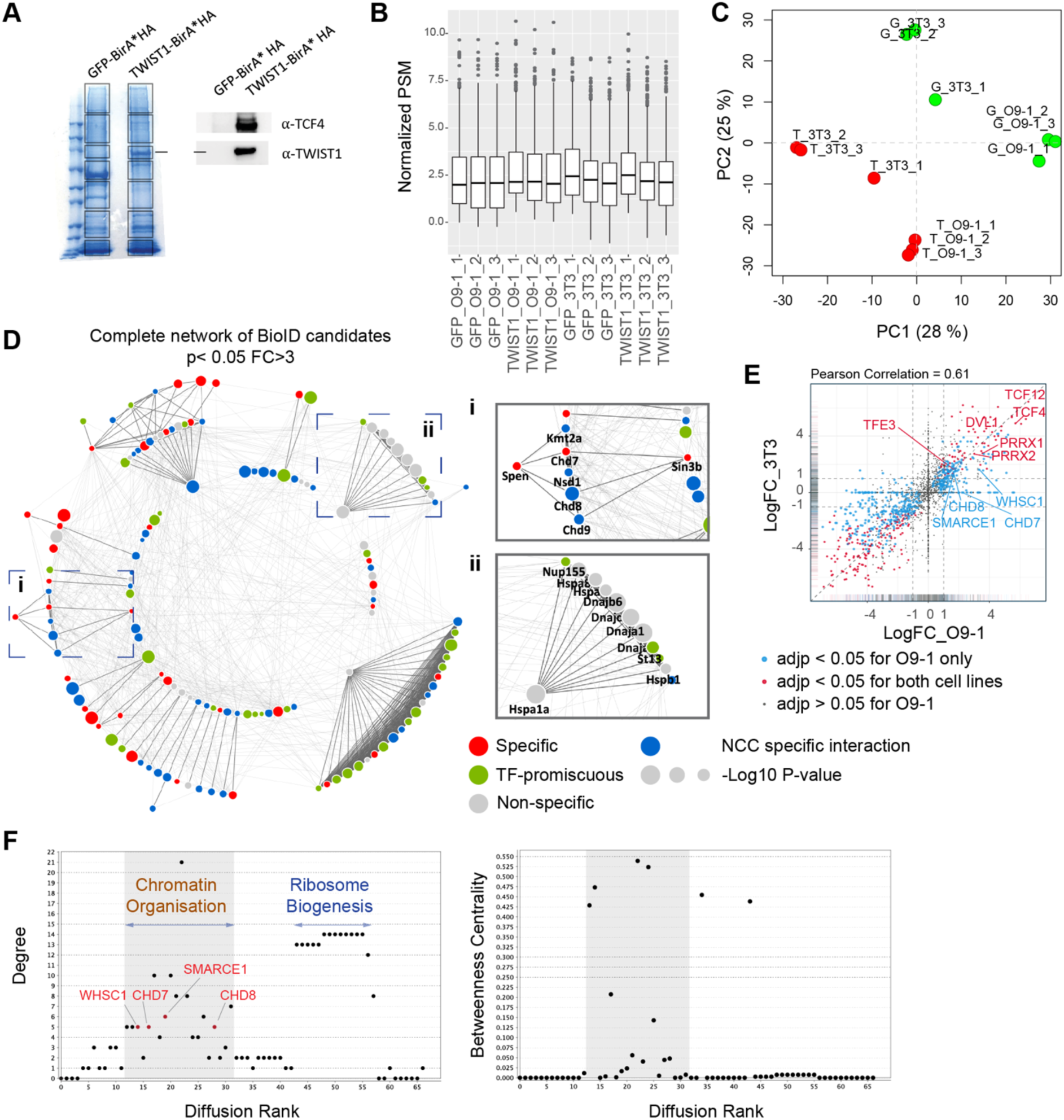
Identification of core NCC regulators within the TWIST1-CRM. **A.** Profile of streptavidin-purified proteins in GFP-BirA* and TWIST1-BirA*-expressing 3T3 cells visualized by Coomassie staining (left panel). Box: Gel bands sampled for mass spectrometry analysis. Expression of the TWIST1-BirA*HA encoded by the transgene and TCF4, a known TWIST1 interactor, by Western blot analysis of the streptavidin-beads purified proteins (right panel). **B.** Mean peptide spectrum match (PSM) across samples, normalized by total PSM of the peptide library. **C.** PCA plot of normalized PSM data. Green dots, GFP-BirA* (G) groups; Red dots, TWIST1-BirA*(T) groups. **D.** Complete network of 140 BioID candidates (P < 0.05; Fold-change > 3; PSM# > 2) interacting physically with TWIST1 in the O9-1 neural crest stem cells. Functional interactions (edges) of these candidates based on prior evidences of co-expression, protein-protein interaction, evolutionary conservation and text mining were retrieved from STRING database (Szklarczyk *et al*., 2015). Medium confidence (combined score > 0.4) was used as the cut-off for interactions. The MCL algorithm was used to generate protein interaction hubs with strongest connection (dark edges). Result from previous protein interaction survey of 56 TFs (Li *et al*., 2015) was referenced to annotate putative specific (red), non-specific (grey) or promiscuous TF interactors (green) among the BioID candidates. Blue nodes are putative specific TWIST1 partners not annotated in Li *et al*. study. Node size = −Log10 (P-value). **i** & **ii.** Example clusters**. E.** Pairwise correlation of TWIST1 BioID data from O9-1 NCCs (x-axis) versus 3T3 fibroblasts (y-axis). Each data point represents one protein, plotted with their log2 fold change of PSM in TWIST1-BirA* versus GFP group (Log2FC). Point density were represented by rug plot next to the axis. Adjusted P-values (adjp) were generated with EdgeR package using negative binomial model: blue, adjp < 0.05 (significant) for O9-1 but not 3T3, red, adjp < 0.05 for both cell lines, black, adjp > 0.05 for O9-1. **F.** Plots generated by NetworkAnalyzer (Assenov *et al*., 2008) for Diffusion Rank of nodes against (**i**) Degree of connection (number of edges), (**ii**) Between-ness centrality, a measures of how fast information spreads to other nodes. Cluster where peaks of highly connected nodes were labelled in (i). Putative neural crest disease factors are likely to arise in the shaded region.

**Figure S3.**
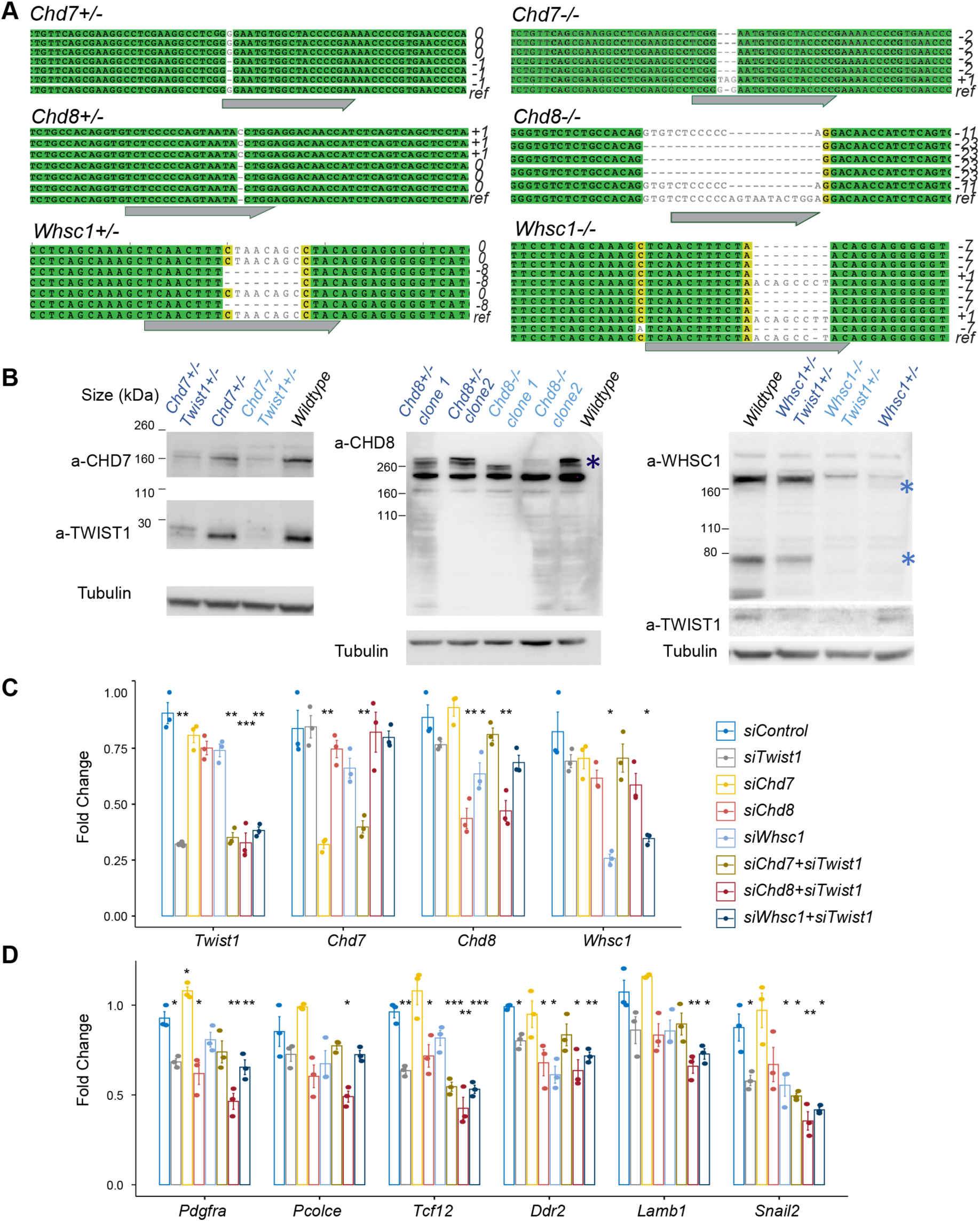
Characterization of CRISPR knockout clones and siRNA knockdown efficiency. **A**. Genotyping result of CRISPR targeted locus in clones with frame-shift mutations. ref, reference/wildtype; Grey arrow, guide RNA targeted site. **B.** Western blot analysis of protein using corresponding antibodies. Expression of Twist1-chromatin regulators was induced by neurogenic differentiation treatment (day 3). Predicted protein sizes were marked by * in WHSC1 and CHD8 blots. qPCR analysis of O9-1 NCCs after 24-hour siRNA treatment (see Methods). **C**, **D**. qPCR results for siRNA targeted genes and EMT markers in the knockdown groups. qPCR signals were normalized against average expression of three housekeeping genes (*Gapdh, Tbp, Actb*) and displayed as fold change +/− SE against control for each treatment. P-values generated using one-way ANOVA. *P < 0.05, **P < 0.01, ***P < 0.001. ns, not significant.

**Figure S4.**
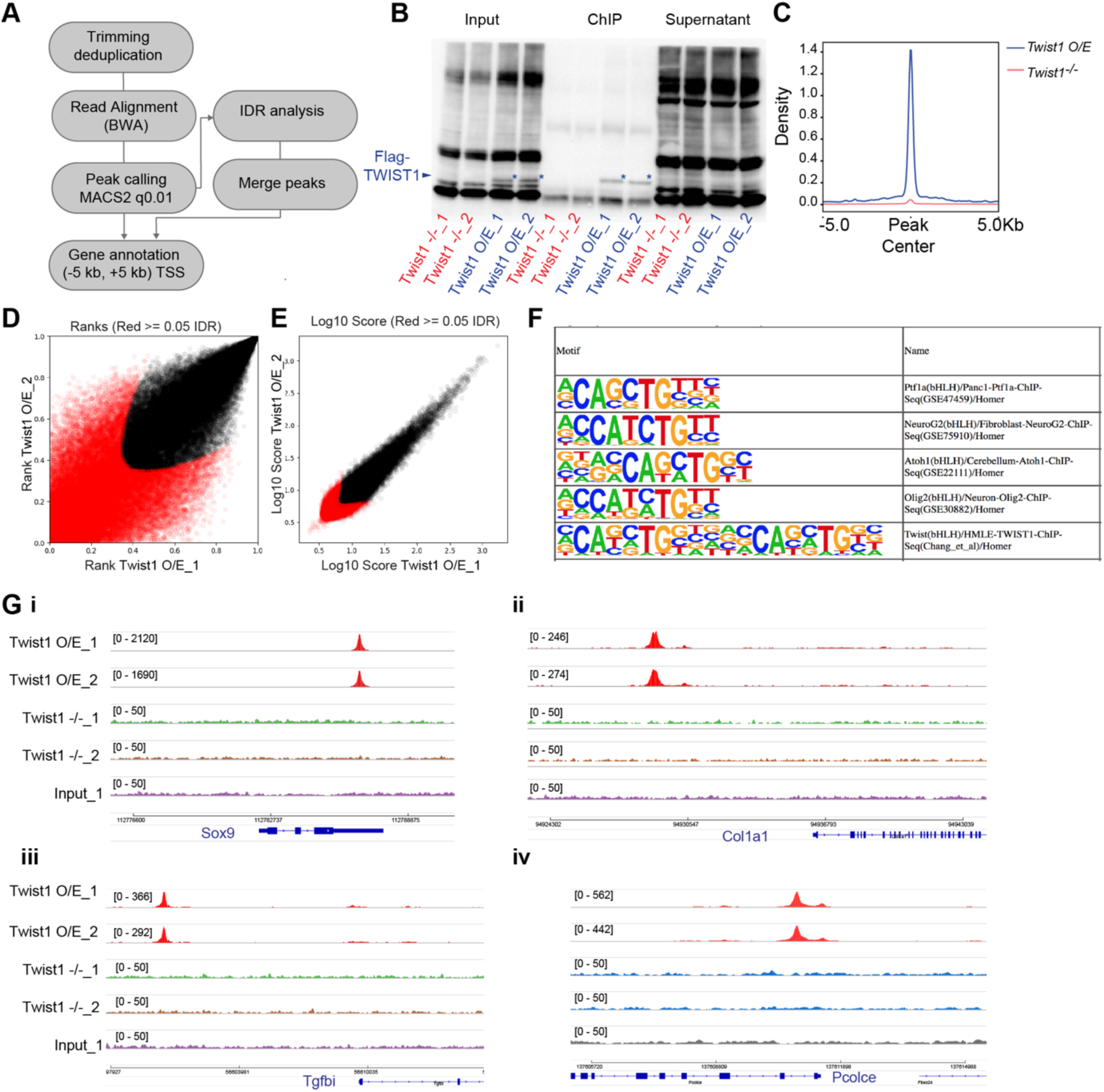
TWIST1 ChIP-seq experiment in ESC-derived neuroepithelial cells. **A.** ChIP-seq data analysis pipeline adapted from ENCODE project (Encode, 2012). **B.** Quality control of chromatin immunoprecipitation specificity. * TWIST1 protein band. **C.** ChIP-seq density profile (rpkm) on mouse genome. **D, E.** IDR analysis showing peaks with significant correlation between replicate experiments (black dots). **F.** Motif enriched in TWIST1-bound chromatin regions. **G.** IGV track showing specific DNA amplification at validated TWIST1 target regions in *Twist1 O/E* but not *Twist1-null* or input ChIP samples.

**Figure S5.**
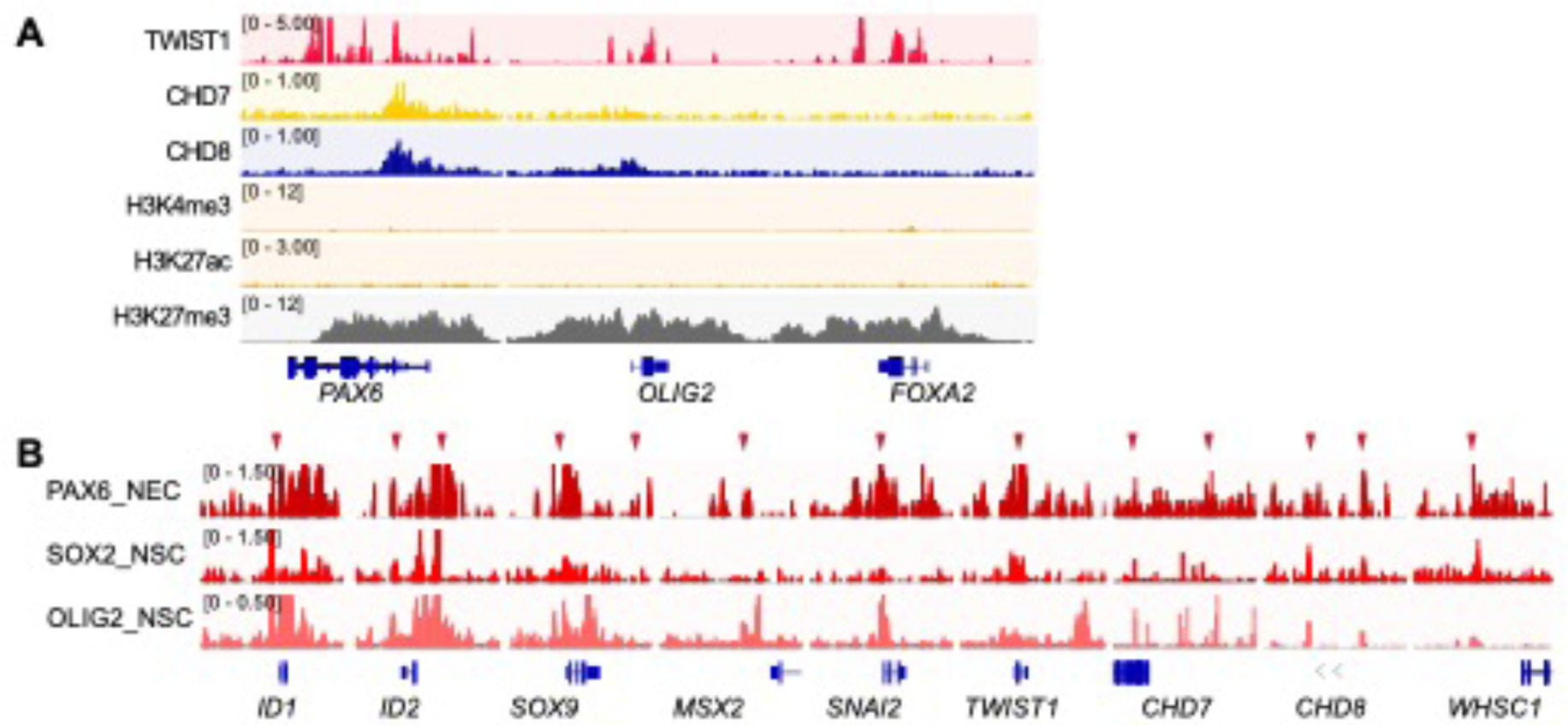
Chromatin binding of TWIST1 and interactors and NSC transcription factors. A. ChIP-seq signal of core TWIST1-chromatin regulators and H3K27me3 histone mark at NSC transcription factors, demonstrating repressed chromatin state. Genes diagrams are indicated (bottom row). B. ChIP-seq signal overlaps (red arrows) for NSC transcription factors at gene locus of NCC specification and NCC-CRM factors. Genes diagrams are indicated (bottom row). Data for NSC TFs (Hikichi *et al*., 2013; Mistri *et al*., 2015; Kutejova *et al*., 2016) were obtained from the Cistrome database (Mei *et al*., 2017).

**Figure S6.**
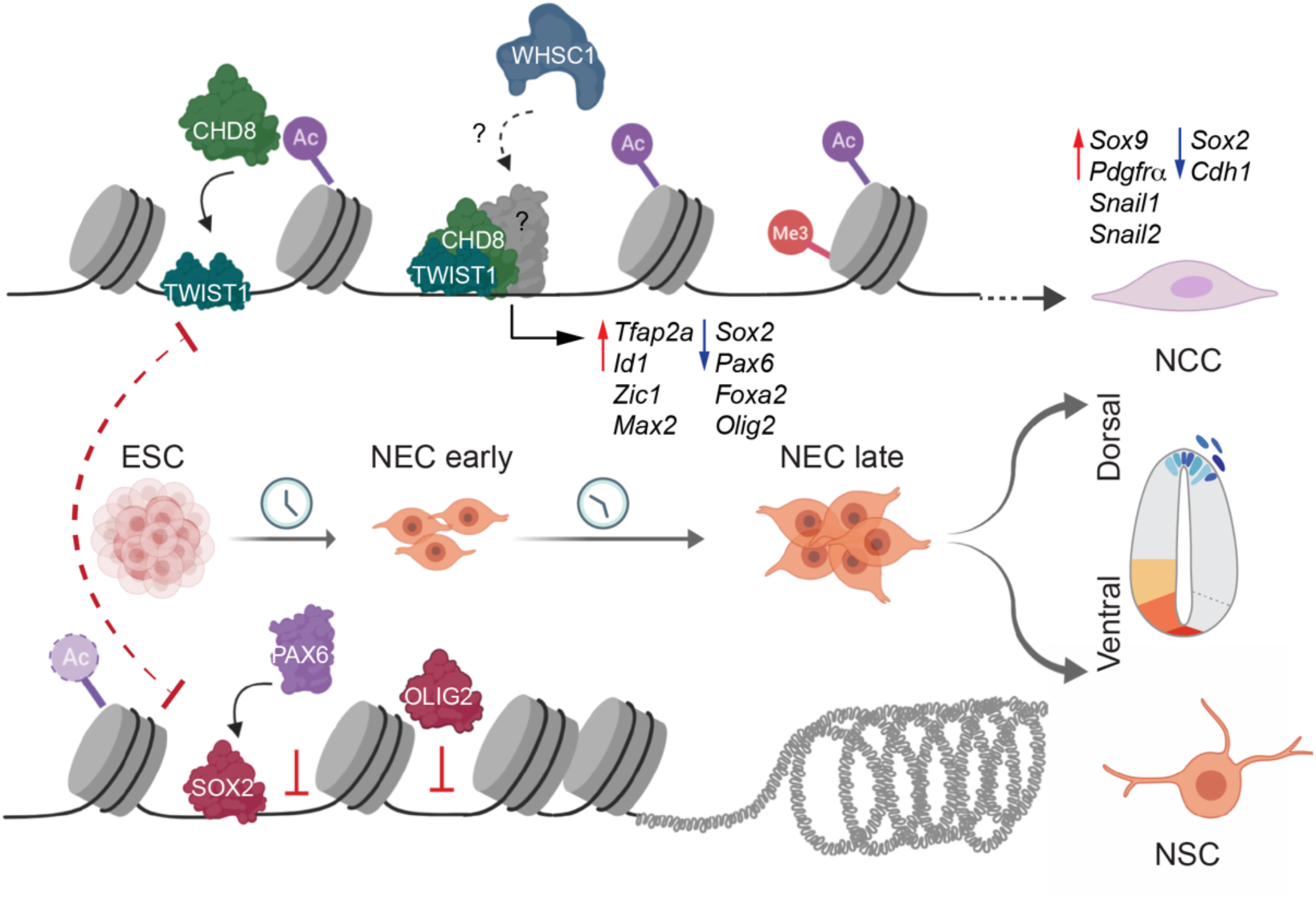
Molecular model of the NCC vs NSC fate decision in neuroepithelium. During NCC specification and dorsal-ventral neural tube axis formation, TWIST1 initiates the assembly of chromatin regulators at the regulatory elements of genes that promote NCC identity and EMT, while repress the expression of NSC TFs. Meanwhile, the NSC TFs: SOX2, PAX6 and OLIG2, competitively occupies these regions to restrict transcriptional activities of the NCC program and enhance the bias towards NSC fates.

## Supplemental Tables

**Table S1** BioID EdgeR test result table

**Table S2** TWIST1 protein interaction module and Gene Ontology analysis

**Table S3** BioID candidates selected for validation

**Table S4** Integrative analysis of ChIP datasets

**Table S5** BioMark reporter card setup

**Table S6** Key resources

**Table S3.**
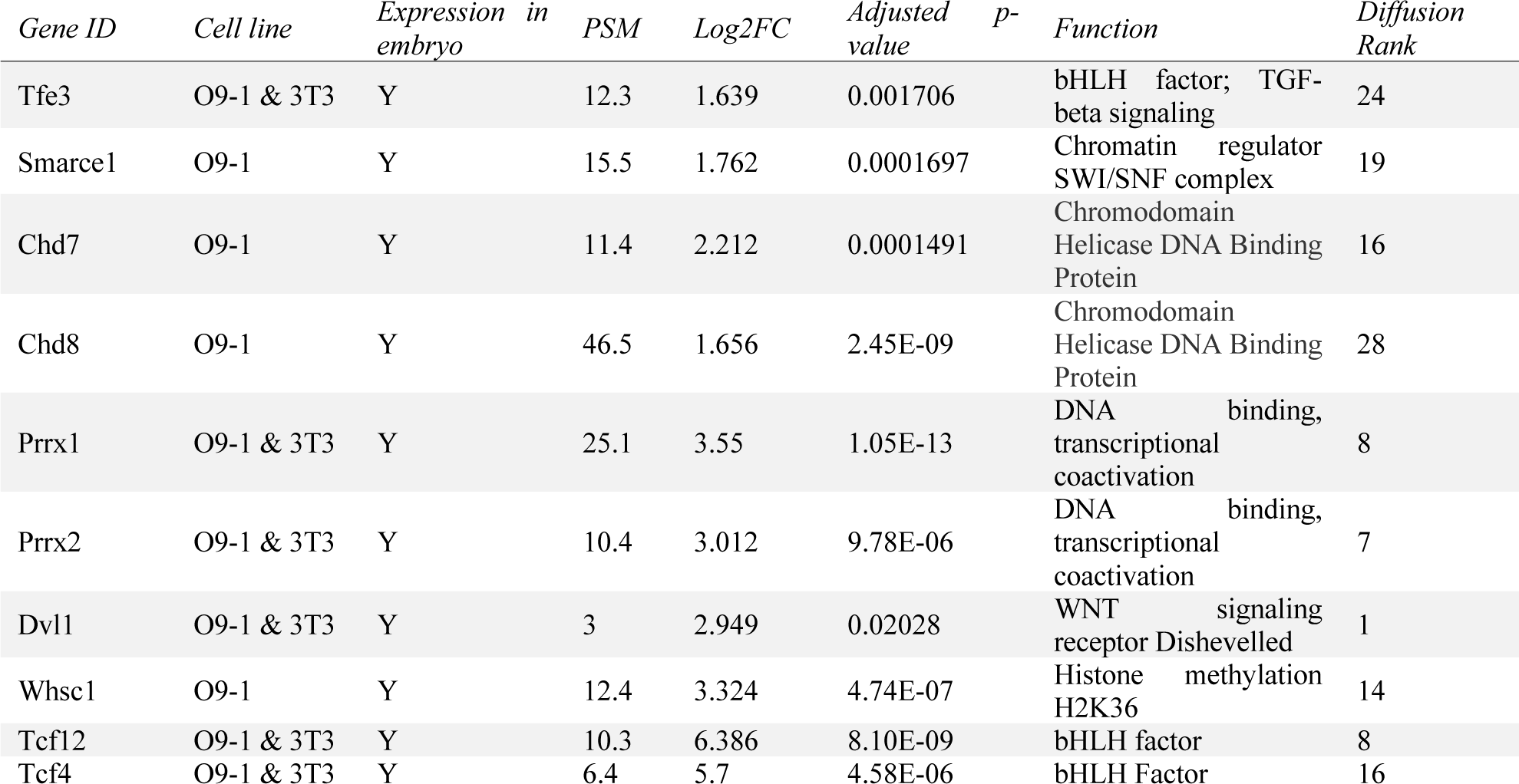
**Information on BioID candidates selected for validation** Cell line of origin of the candidate is listed. Expression data of the embryonic head was from published study (Fan *et al*., 2016). Log2 FC = log2 transformed PSM fold-change between TWIST1-BirA*HA and GFP transfected O9-1 cells. Adjusted p-value was computed from dataset from O9-1 cell line, generated by the likelihood ratio test corrected by the Benjamini & Hochberg method in EdgeR (Robinson *et al*., 2010). The rank of candidates in heat diffusion from genes associated with human and mouse facial malformation are listed.

